# Targeting *Fusobacterium nucleatum* through Chemical Modifications of Host-Derived Transfer RNA Fragments

**DOI:** 10.1101/2022.09.29.510195

**Authors:** Mengdi Yang, Pu-Ting Dong, Lujia Cen, Wenyuan Shi, Xuesong He, Jiahe Li

## Abstract

Host mucosal barriers possess an arsenal of defense molecules to maintain mucosal health. In addition to well-established defense molecules such as antimicrobial peptides and immunoglobulins, a subset of extracellular host-derived small RNAs (sRNAs) also exhibits antimicrobial functions in a cross-kingdom fashion. We recently uncovered the sRNA-mediated crosstalk between human normal oral keratinocytes and *Fusobacterium nucleatum (Fn)*, an opportunistic oral pathobiont with increasing implications in extra-oral diseases. Notably, when challenged with *Fn*, oral keratinocytes released *Fn*-targeting tRNA-derived sRNAs (tsRNAs), an emerging class of noncoding sRNAs with diverse functions in gene regulation. Additionally, synthetic mimics of the *Fn*-targeting tsRNAs exhibited highly selective antimicrobial activity against *Fn*. However, excess synthetic tsRNAs (in the micromolar range) were required to achieve growth inhibition, which limits their potential as antimicrobials. Here, we chemically modify nucleotides of the anti-*Fn* tsRNAs, termed MOD-tsRNAs, and demonstrate their species- and sequence-specific inhibition in the nanomolar range in various *Fn* type strains and clinical tumor isolates. In contrast, the same MOD-tsRNAs do not inhibit two representative oral bacteria, *Porphoromonas gingivalis* (*Pg*) and *Streptococcus mitis* (*Sm*). Additionally, MOD-tsRNAs are internalized by different *Fn* strains while exhibiting minimal uptake by *Pg* and *Sm*. Further RNA sequencing and affinity pull-down assays implicate MOD-tsRNAs as potential ribosome-targeting antimicrobials against *Fn*. Taken together, our work provides a framework to target opportunistic pathobionts through co-opting host-derived extracellular tsRNAs, whose potential applications may have been limited by their intrinsic instability as well as our limited understanding of the inhibition mechanism.

## INTRODUCTION

Host mucosal surfaces are highly specialized and possess a complex array of innate and adaptive immunity^[1, 2]^. They provide the first line of protection against infectious agents by initiating protective responses to potential pathogens. Furthermore, the symbiotic relationship of the hundreds of microbial species with the host requires a fine-tuned response at the mucosal surface that prevents overgrowth of opportunistic pathogens, while sparing beneficial microbes^[2]^. As a result, multiple innate and adaptive mechanisms involving antimicrobial peptides, complement and immunoglobulins have evolved to maintain the delicate balance between the host and associated microbiomes^[3-5]^. In addition to these well-established systems, recent studies have begun to shed light on the roles of host-derived small RNAs (sRNAs) that contribute to the maintenance of host-microbial homeostasis through cross-kingdom gene modulation^[6, 7]^. For instance, eukaryotic cells can secrete certain sRNAs (*e*.*g*., miRNAs) into extracellular environments, either encased in extracellular vehicles (EVs) or in an EV-free mode. These sRNAs can then act on distantly related organisms and exert regulatory functions in a cross-kingdom fashion^[8-10]^. While mounting evidence has demonstrated that plants and vertebrate animals exploit extracellular miRNAs as a defense mechanism in plant-pathogen and host-gut microbiota interactions^[11-13]^, we recently identified an emerging class of host-derived sRNA, named transfer RNA-derived small RNA (tsRNA) with implications in the host-bacteria interactions^[14]^. While tsRNAs were originally identified to regulate gene expression inside eukaryotes in a cell autonomous manner^[15]^, accumulating evidence indicates that distinct tsRNA species are produced and secreted by host cells under various physiological and pathological conditions, some of which have been proposed to serve as disease biomarkers^[16]^. In addition to these established roles, host-derived tsRNAs were recently implicated in the cross-kingdom interactions between human oral epithelial cells and *Fusobacterium nucleatum* (hereinafter *Fn*)^[14]^. Specifically, *Fn* is a key opportunistic oral pathobiont and has garnered renewed attention in recent years. Specifically, in addition to being a bridging bacterium of dental plaque and its roles in periodontitis, it has been postulated that *Fn* is disseminated systemically from oral cavity to other organs contributing to extra-oral diseases such as adverse pregnancy outcomes, colorectal cancer, Alzheimer’s disease and various other diseases^[17-19]^. To investigate a possible mechanism to regulate *Fn* by the host, we previously identified a total of 68 distinct human tsRNAs using small RNA sequencing in cell-free saliva from four healthy human subjects, among which two host-derived tsRNAs, termed tsRNA-000794 and tsRNA-020498, bear high sequence homology with tRNAs in *Fn*. Interestingly, immortalized human oral epithelial cells released tsRNA-000794 and tsRNA-020498 in response to *Fn* challenge in an *in vitro* bacteria-host coculture model^[14]^. Furthermore, we made two key observations: (1) chemically synthesized RNA oligos that mimic the exact compositions of naturally occurring tsRNA-000794 and tsRNA-020498, inhibited the growth of *Fn* ATCC 23726, but not *Streptococcus mitis* ATCC 6249 (*Sm*), a health-associated gram-positive oral bacterium; and (2) these two tsRNA mimics inhibited *Fn* in a sequence-dependent manner. Of note, a scrambled RNA control or a prevalent human salivary piRNA (piRNA-0016792), which bears no sequence homology to *Fn* tRNAs, failed to inhibit *Fn*. These findings not only uncovered another layer of defense mechanism by which the host modulates its interaction with associated microbes, but also suggested a new research avenue to repurpose host sRNAs for tool development and translational applications by achieving targeted depletion of disease-associated bacteria. However, despite the observed specificities at the sequence and species levels, excess synthetic tsRNA mimics (in the micromolar range) were required to inhibit *Fn*. This poses a formidable challenge to co-opting host-derived tsRNAs as a potential antimicrobial agent targeting *Fn*.

In the present work, to address the limitations, we drew on the power of rapid advances in chemically modified RNAs towards development of powerful genetic tools and therapeutic reagents, including FDA-approved small interfering RNA-based drugs^[20]^, messenger RNA vaccines^[21]^, and guide RNA for genome editing by Clustered Regularly Interspaced Short Palindromic Repeats (CRISPR)^[22, 23]^. Specifically, we adapted a similar chemical modification strategy to generate two modified tsRNAs (referred to as MOD(OMe)-tsRNAs in this study) matching the exact sequences of tsRNA-000794 and tsRNA-020498 (Table 1), which resulted in nearly 1000-fold increase in the efficacy of inhibiting *Fn* growth while preserving the sequence and species specificities. We then demonstrated the uptake of MOD(OMe)-tsRNAs by multiple *Fn* strains of oral and gastrointestinal origins. Furthermore, RNA-seq analysis, tsRNA pull-down assay and Raman spectroscopy provided evidence that MOD(OMe)-tsRNAs likely inhibit *Fn* through a mechanism reminiscent of the mode of action for ribosome-targeting antimicrobials. Taken together, our work exemplifies interdisciplinary approaches to understanding and harnessing host-derived tsRNAs to target opportunistic oral pathobionts in a sequence- and species-specific fashion. Moreover, the study provides insight into the potential roles of host-derived tsRNAs in regulating bacterial physiology and host-microbial interactions.

**Table 1.**
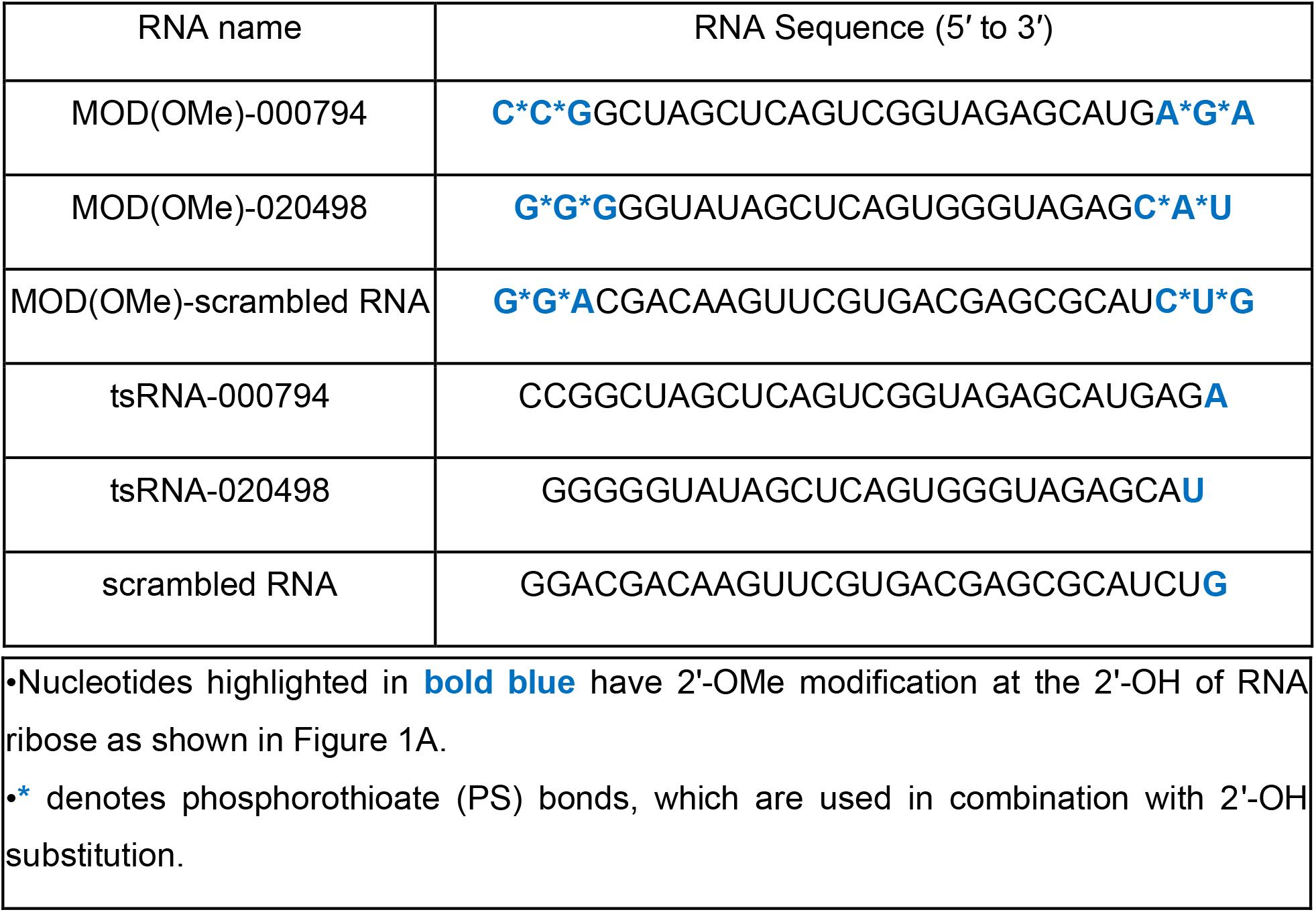
List of chemically modified and naturally occurring tsRNAs

## RESULTS

### Chemical modifications of host-derived tsRNAs enhance the inhibitory efficacy while maintaining the sequence and species specificities

Host derived extracellular tsRNAs are typically encapsulated in extracellular vesicles or associated with proteins to confer tsRNAs stability in human saliva. In our earlier work, directly adding synthetic mimics of naturally occurring tsRNAs inhibited the growth of *Fn* albeit with a low efficacy (the half maximal inhibitory concentration (IC50) is ∼ 50 µM), likely due to the susceptibility of naked RNA to nuclease-mediated degradation in bacterial culture. To increase tsRNAs stability, we explored two common RNA modifications (Fig. 1*A*): First, a methyl group was added to the 2’ hydroxyl (2’-O-methylation) of the ribose moiety of the three terminal nucleosides at both 5’ and 3’ ends^[23]^. Of note, 2’-O-methylation is prevalent in both prokaryotes and eukaryotes as a key post-transcriptional RNA modification for noncoding RNA species, among which piRNAs and miRNA bear a 2’-O-methylated nucleotide at the 3’ end^[24-26]^. Second, two phosphodiester linkages were replaced with phosphorothioate (PS) bonds at both 5’ and 3’ termini, in which a non-bridging oxygen in the phosphodiester bond was substituted with a sulfur atom. While these two modification strategies have been widely used to confer nuclease resistance on guide RNA and siRNAs for genome editing and RNA interference^[20, 23]^, it has yet to be confirmed whether 2’-O-methylation and PS modifications can also improve the stability and *Fn*-inhibiting efficacy of the naturally occurring tsRNAs that we previously identified. For the rest of this study, we will name MOD(OMe)-000794, MOD(OMe)-020498 and MOD(OMe)-scrambled to specify 2’-O-methylation and PS bond for tsRNA-000794, tsRNA-020498 and the non-targeting scrambled RNA control, respectively. Using a stem-loop reverse transcription polymerase chain reaction (RT-PCR) assay, we showed that MOD(OMe)-000794, MOD(OMe)- 020498 and MOD(OMe)-scrambled indeed displayed markedly improved stability by approximately 100 times compared to that of the naturally occurring ones when incubated in bacterial culture media (Fig. 1*B*, Fig. S1).

**Fig. 1.**
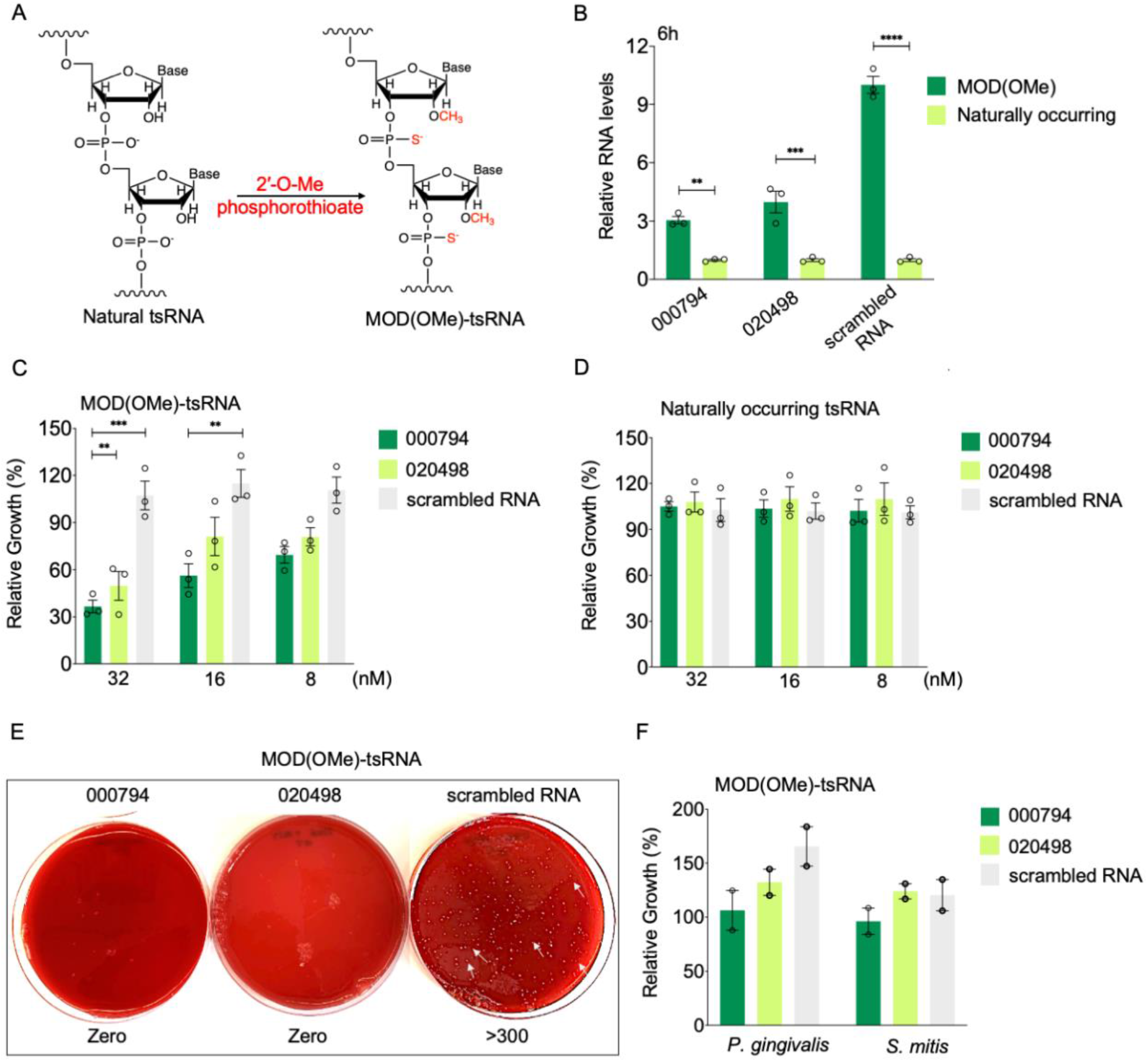
Chemically modified tsRNAs confers superior growth inhibition of *Fn* in sequence-, and species-dependent manner. (*A*) Incorporation of 2’-O-methyl phosphorothioate linkage in tsRNAs, referred to as MOD(OMe)-000794, MOD(OMe)-020498 and MOD(OMe)-scrambled RNA. (*B*) Improvement of tsRNA stability by chemical modifications in comparison to synthetic mimics of naturally occurring counterparts. Individual tsRNAs (1 nM) were incubated with Columbia broth anaerobically at 37 °C for 6 h, and intact tsRNAs were measured via quantitative real time PCR using a stem-loop method. Results were normalized to the initial levels of tsRNAs. (*C*) Enhanced growth inhibition of *Fn* ATCC 23726 by MOD(OMe)-000794 and MOD(OMe)-020498 at the nanomolar concentration ranges, but not MOD(OMe)-scrambled RNA. Results were representative of three biological replicates. (*D*) Naturally occurring tsRNAs failed to inhibit *Fn* ATCC 23726 at same concentrations. (*E*) Representative images showing the colony formation of *Fn* ATCC 23726 on agar plates after treatment with 500 nM MOD(OMe)-000794, MOD(OMe)-020498 or MOD(OMe)-scrambled RNA for 24 h. Bacteria were washed once and recovered on nonselective Columbia broth sheep blood agar plates. (*F*) MOD(OMe)-000794 and MOD(OMe)-020498 at 512 nM exhibited no growth inhibition in two oral bacteria, a gram-negative bacterium, *Porphyromonas gingivalis* ATCC 33277 (*P. gingivalis*) and a gram-positive bacterium, *Streptococcus mitis* ATCC 6249 (*S. mitis*). Results were representative of two biological replicates. Data were analyzed by the two-way ANOVA followed by Dunnett’s Bonferroni multiple comparison tests. *p < 0.05, **p < 0.01, ***p < 0.001, ****p < 0.0001.

Having validated the enhanced stability of modified tsRNAs over their naturally occurring counterparts, we further assessed whether MOD-tsRNAs exhibit increased efficacy of inhibition against *Fn*. As shown in Fig. 1*C*, MOD(OMe)-000794 and MOD(OMe)-020498 inhibited the growth of *Fn* ATCC 23726 in a dose-dependent manner, achieving an IC50 in the range of 16-32 nM. In comparison, at the same concentrations, naturally occurring tsRNAs did not affect the growth of *Fn* ATCC 23726 (Fig. 1*D*). In addition, MOD(OMe)- scrambled RNA control did not inhibit *Fn* ATCC 23726 even at 500 nM, indicating that the observed inhibition was dependent on the specific tsRNA sequence rather than chemical modifications (Fig. S2). Meanwhile, MOD(OMe)-000794 and MOD(OMe)-020498, but not MOD(OMe)-scrambled RNA control, also inhibited the growth of two additional *Fn* type strains, *Fn* ATCC 25586 and 10953 (Fig. S3), albeit with slightly reduced efficacy.

Considering a ∼1000-fold increase in growth inhibition efficacy by MOD-tsRNAs compared to their naturally occurring tsRNA counterparts, we further asked whether *Fn* were able to regrow after the treatment with MOD-tsRNAs. To this end, we treated *Fn* ATCC 23726 with 500 nM MOD(OMe)-000794, MOD(OMe)-020498, and MOD(OMe)- scrambled RNA control, respectively for 24 h, and recovered bacteria on non-selective Columbia broth blood agar to enumerate the colony-forming unit (CFU). Notably, 24 h treatment with MOD(OMe)-000794 or MOD(OMe)-020498 resulted in complete killing of *Fn* ATCC 23726 cells (Fig. 1*E*), which was approximately eight orders of magnitude of reduction in the CFU compared to that of MOD(OMe)-scrambled RNA control. Decrease in the CFU was further reproduced in *Fn* ATCC 25586 (Fig. S4), after 24 h treatment with MOD(OMe)-000794 or MOD(OMe)-020498 but not MOD(OMe)-scrambled RNA control. These findings suggested highly potent killing of *Fn* by MOD-tsRNAs.

In addition to the sequence specificity, we tested whether MOD-tsRNAs displayed *Fn*- specific growth inhibition. To this end, we challenged two oral bacteria including Gram- negative *Porphyromonas gingivalis* ATCC 33277 (*Pg*), and Gram-positive *Streptococcus mitis* ATCC 6249 (*Sm*) (Fig. 1*F*), as well as *E. coli* K-12 (Fig. S5) with MOD(OMe)-000794, MOD(OMe)-020498 and MOD(OMe)-scrambled RNA control. Indeed, none of these three bacteria was inhibited by MOD(OMe)-tsRNAs at a concentration of 512 nM (> 10- fold higher than the IC50 of MOD-tsRNAs for *Fn*). Since the two naturally occurring tsRNAs were originally identified in human saliva and can also be secreted by human oral epithelial cells^[14]^, we further tested whether chemically modified tsRNAs affect the viability of host cells. As shown in Fig. S6, 512 nM of MOD(OMe)-000794 or -020498 did not affect the proliferation of immortalized human oral epithelial cells in comparison to the untreated or MOD(OMe)-scrambled RNA control.

Motivated by the enhanced efficacy of tsRNA through partial chemical modifications at the three terminal nucleotides, we further explored and compared fully modified tsRNAs for their efficacy in inhibiting *Fn* growth to naturally occurring and partially modified sequences (Fig. S7*A*). Intriguingly, full modifications of RNA backbone completely abolished the efficacy (Fig. S7*B*), which underscores the degrees of modifications in dictating the anti-*Fn* properties of tsRNAs. These findings are in line with prior studies in RNA interference^[27]^ and CRISPR genome editing^[22, 23]^, which demonstrated that extensive chemical modifications compromised the activities of siRNA and guide RNA likely through altering the secondary structure of RNA, affecting interaction between RNA and target proteins, or inducing nonspecific interactions with cellular components. Taken together, our findings highlighted the importance of the degrees of RNA modifications towards enhanced stability and efficacy of tsRNAs.

### Chemically modified tsRNAs inhibit *Fn* clinical tumor isolates

*Fn* has garnered increasing attention due to its implications in colorectal cancer (CRC)^[28, 29]^. In addition to three type strains (*Fn* ATCC 23726, 25586 and 10953), we further tested *Fn* clinical tumor isolates (CTI) from CRC by challenging them with the same MOD- tsRNAs. We first chose CTI-2 and CTI-7 due to their distinct adhesion characteristics to CRC cells. Specifically, the interaction between D-galactose-b(1-3)-N-acetyl-D- galactosamine (Gal-GalNAc) from cancer cells and a surface protein, Fap2, from *Fn* has been shown to promote the enrichment of fusobacteria in CRC. While CTI-2 expresses Fap2 that mediates the adhesion of *Fn* to colon cancer epithelial cells overexpressing Gal-GalNAc, CTI-7 is *fap2*-deficient and does not efficiently attach to cancer cells^[30]^. Unlike type strains, however, the colon isolates grow at a slower rate under anaerobic conditions, and tend to aggregate to form nonuniform cultures, rendering it less accurate to quantify the growth rates by measuring the absorbance within a short time of incubation. To circumvent this issue, we chose a cell viability dye, SYTOX Green, which emits an intense fluorescence in necrotic bacteria through binding to nucleic acids. We reasoned that the SYTOX Green dye offers an alternative yet more sensitive way to detect the response of *Fn* isolates than that of absorbance measurement. Consistent with the findings in *Fn* ATCC 23726, 25586 and 10953, overnight treatment with MOD(OMe)- 000794 and -020498 resulted in significant cell death in CTI-2 and -7, compared to that of MOD(OMe)-scrambled RNA control (Fig. 2*A* and *B*).

**Fig. 2.**
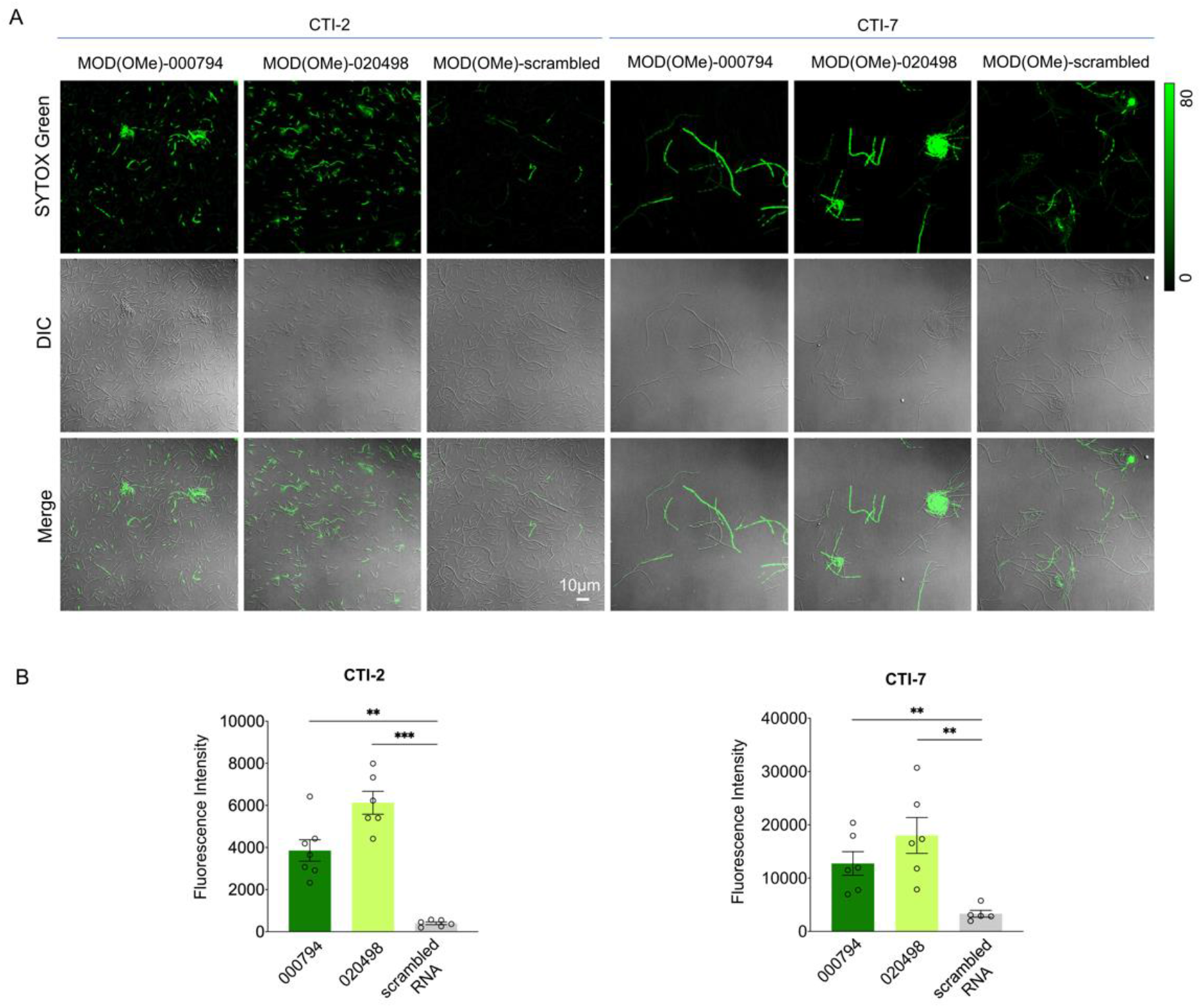
Fluorescence microscopy of SYTOX Green stain-labeled *Fn*. (*A*) *Fn* CTI-2 and CTI-7 were treated with 500 nM MOD(OMe)-000794, MOD(OMe)-020498, and MOD(OMe)-scrambled for 48h and 72h respectively. SYTOX green was added to PBS resuspended bacterial culture and incubated for 30mins. (*B*) SYTOX Green quantification was achieved by calculating the ratio between raw integrated fluorescence intensity and the area of all the *Fn*. Data were analyzed by the Student unpaired t-test: *p < 0.0332, **p < 0.0021, ***p < 0.0002, ****p < 0.0001.

### Internalization of *Fn*-targeting tsRNAs by *Fn* but not *Sm* or *Pg*

Prior studies in host-gut microbiome and plant-fungi interactions demonstrated that host-derived sRNAs can enter bacteria and fungi to modulate their physiology^[11-13]^. Given the ability of two tsRNAs in inhibiting *Fn*, we next asked whether the *Fn*-targeting tsRNAs also entered *Fn* to exert the growth inhibition. We fluorescently labeled the tsRNAs with Cy3 at the 3’ end and treated *Fn* ATCC 23726 with tsRNA-000794-Cy3, tsRNA-020498- Cy3, or the scrambled-RNA-Cy3 (Fig. 3*A*). After overnight incubation, we examined the localization of fluorescently labeled tsRNAs in *Fn* ATCC 23726 through the Airyscan confocal microscopy, an optical technique that allows us to detect morphological features at a higher resolution than that of conventional microscopy^[31]^. As shown in Fig. 3*B*, the majority of tsRNA-000794-Cy3 accumulated in the cytoplasm of *Fn*, while there was much lower uptake for the scrambled RNA. A similar uptake profile was observed in *Fn* ATCC 25586 (Fig. S8), suggesting that the two *Fn* strains likely shared the same mechanism in internalizing tsRNAs. Further quantification indicated that tsRNA-000794-Cy3 exhibited higher intracellular accumulation than tsRNA-020498-Cy3 in *Fn* ATCC 23736 (Fig. 3*A*), which agreed with the slightly higher growth inhibition of *Fn* 23726 by MOD(OMe)-000794 than that of MOD(OMe)-020498 (Fig. 1*C*). For these reasons, we subsequently focused on characterizing tsRNA-000794-Cy3 in the two clinical tumor isolates. As shown in Fig. S9, Cy3-labeled tsRNAs were also localized in the cytoplasm of clinical tumor isolates after 24 h incubation, which further suggested that MOD(OMe)-tsRNAs entered *Fn*, regardless of their origins, to mediate the growth inhibition (Figs. 1*C* and 2*B*). We next tested whether the uptake of tsRNA is dependent on the energy metabolism in *Fn* by using sodium azide, a common metabolic inhibitor^[32]^. The internalization of tsRNA- 000794-Cy3 was markedly reduced by 0.6 mM sodium azide (Fig. 3*C*). Of note, this concentration, which was 100-fold lower than that used for inhibition of the energy metabolism in *Fn* ^[32]^, did not significantly inhibit the growth of *Fn* ATCC 23726 (Fig. S10), suggesting that the uptake of tsRNAs is dependent on an active transport mechanism rather than passive diffusion. To confirm that the observed intake was not due to specific properties of the linked dye molecule, we next substituted Cy3 with Alexa-488, a structurally and spectrally different dye molecule. As expected, we consistently observed enhanced uptake of tsRNA-000794-Alexa488 and tsRNA-020498-Alexa488 compared to the scrambled RNA-Alexa488 (Fig. S11*A*). Additionally, two Alexa-488 conjugated tsRNAs but not the scrambled RNA could inhibit the growth of *Fn* ATCC 23726 (Fig. S11*B*). Taken together, our results showed that (1) tsRNAs can be internalized by *Fn*; and (2) 3’ end labeling of tsRNA with a fluorescent dye did not significantly interfere with its inhibitory effect against *Fn*.

**Fig. 3.**
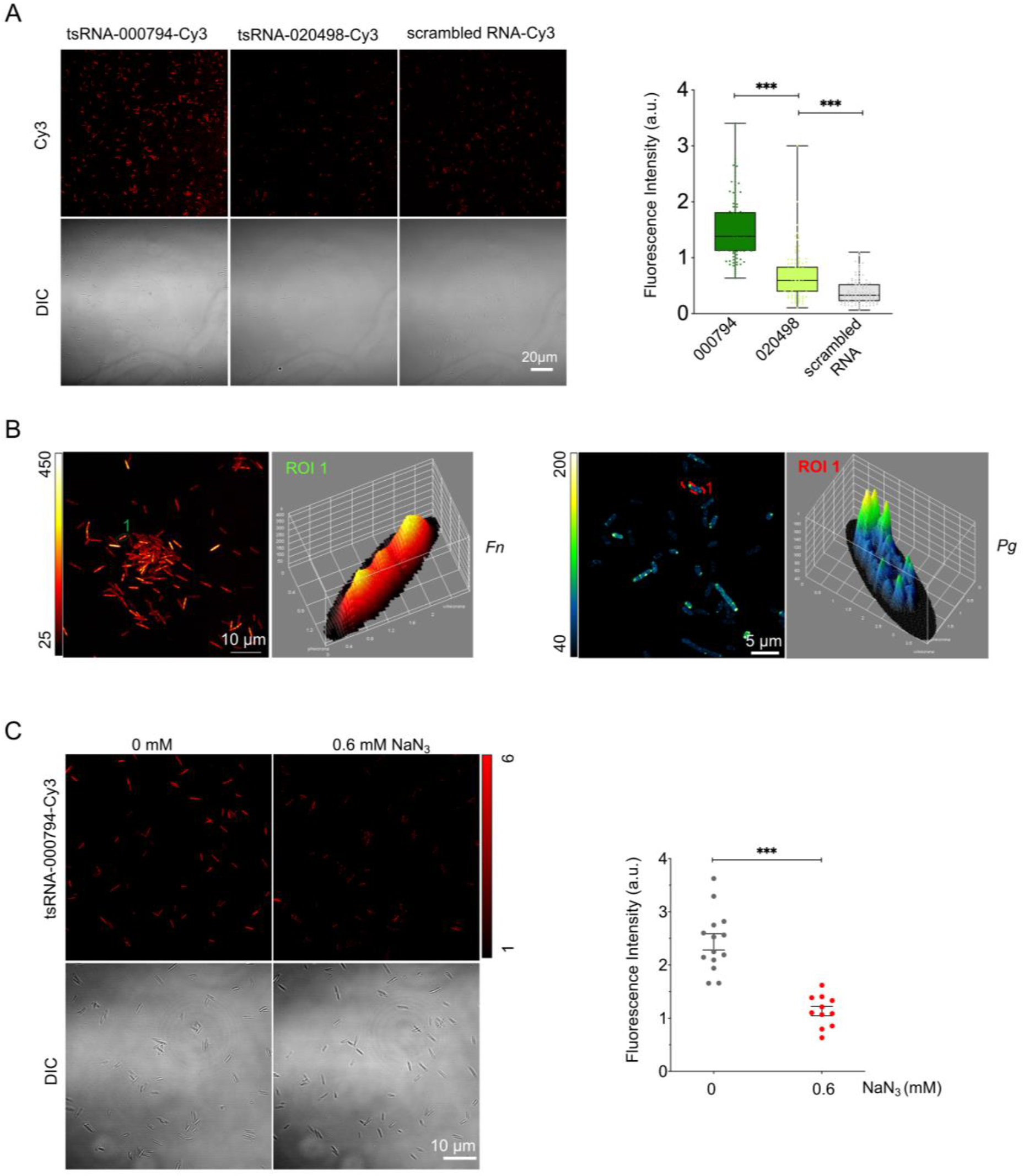
Specific Internalization of host-derived tsRNAs by *Fn*. (*A*) Treatment of *Fn* ATCC 23726 with tsRNA-000794-Cy3, tsRNA-020498-Cy3 or scrambled RNA-Cy3, and subsequent quantification of fluorescence intensity reflecting the abundance of tsRNAs in *Fn*. Box-and-whisker plots: median, horizontal line; box range, percentile 25, 75. Data was determined by Student unpaired t-test (***: p<0.001; **: p<0.01). (*B*) Intracellular accumulation of tsRNA-000794-Cy3 in *Fn* ATCC 23726 compared with periphery localization in *Pg*. The images were acquired by Airyscan confocal microscopy. Individual Cy3-labeled RNA oligos were incubated with indicated bacteria in nutrient rich broth under anaerobic conditions for 16 h, washed with 0.9% saline twice to remove free RNA, and then mounted on glass slides for airyscanning. Images are representative of three biological replicates. (*C*) The uptake of tsRNA-000794-Cy3 in the absence of presence of 0.6 mM sodium azide. Quantification of fluorescence intensity for tsRNA-000794-Cy3 in randomly selected bacteria. Statistical significance was determined by the Student unpaired t-test(***: p<0.001; **: p<0.01).

Since the two MOD-tsRNAs can specifically kill *Fn* but spare other oral bacteria including *Pg* and *Sm*, we further asked whether the selective killing by MOD-tsRNAs against *Fn* can be attributed to different uptake patterns in *Pg* and *Sm*. Intriguingly, in contrast to cytoplasmic accumulation of tsRNA-Cy3 in *Fn* ATCC 23726, *Fn* ATCC 25586 and clinical tumor isolates, the same tsRNA-Cy3 was primarily located at the periphery of *Pg* and *Sm* (Fig. 3*B*, Fig. S12) as evidenced by the Airyscan confocal microscopy imaging. While the differential uptake patterns for tsRNA-Cy3 between *Fn, Pg* and *Sm* may be implicated in the targeted growth inhibition against tsRNAs in *Fn*, it remains unknown how MOD- tsRNAs affected *Fn*.

### Global RNA profiles show MOD-tsRNAs target protein translation

To answer the above question, we next performed bacterial RNA sequencing (RNAseq) to compare transcriptomic differences between MOD(OMe)-000794 and MOD(OMe)- scrambled control treated *Fn* ATCC 23726. It is critical to optimize both concentrations and treatment duration such that the transcriptomics does not merely reflect cell death-associated gene expression changes. Meanwhile we reasoned that if *Fn* ATCC 23726 is treated with lower concentrations or shorter duration than the optimal conditions, there may not be significant transcriptomic changes to infer the targets and functions of MOD- tsRNAs. For these reasons, global transcriptome studies of antimicrobial responses generally use either sub-inhibitory concentrations of the inhibitors of interest for a relatively long duration or analyze transcriptome profiles soon after exposure to a lethal concentration, with each approach having its advantages and disadvantages^[33]^. Here, we opted for RNA-seq at an early time point after treating *Fn* ATCC 23726 with inhibitory doses of MOD(OMe)-000794 or MOD(OMe)-scrambled RNA control and analyzed samples by bacterial RNAseq. Since we focused on the short-term response of *Fn* when exposed to a lethal concentration of MOD-tsRNA, a 10-time higher starting OD (OD_600_=0.2) than the viability assay was used to obtain enough bacteria for RNA extraction. However, it was difficult to rely on absorbance measurement at early time points to optimize the concentrations of MOD-tsRNAs that can induce detectable inhibition in the bacterial samples. To address the issues, the SYTOX green viability dye was employed to monitor differences in cell viability when *Fn* ATCC 23726 were treated with MOD(OMe)-000794 or MOD(OMe)-scrambled at various concentrations within a short treatment period. Of note, approximately ∼10% reduction in cell viability for *Fn* ATCC 23726 was observed under the treatment condition at 500 nM MOD(OMe)-000794 for 5 h (Fig. S13), which corresponds to approximately one round of cell division for *Fn* according to our experiment and the literature^[34, 35]^.

After optimizing the treatment conditions, gene expression levels were compared between MOD(OMe)-000794 (treatment) and MOD-scrambled (control) in three biological replicates. Differentially expressed genes designated as those with a false discovery rate (FDR)-adjusted P-value < 0.001 and absolute fold change >2 are presented by heatmaps (Fig. 4*A*) and volcano plots (Fig. 4*B*). As shown in Table 2, genes associated with putative purine metabolism and hemin uptake pathways were significantly downregulated in the treatment group. In addition, genes related to chaperones, ribosomal proteins, and tRNAs displayed up to 4-fold upregulation, suggesting that treatment with MOD(OMe)-000794 likely affected protein folding and production. We further validated these genes with >2-fold down- and up-regulation by real time PCR in three biological replicates and the results were consistent with those in the RNA-seq data (Fig. 4*D*). To put these genes into a functional context, we performed a pathway enrichment analysis of differentially regulated genes based on the Kyoto Encyclopedia of Genes and Genomes (KEGG) database (Fig. 4*C*). The top five most enriched pathway terms are ribosome, RNA degradation, purine metabolism, pentose phosphate pathway and fatty acid/lipid biosynthesis, which were affected by MOD(OMe)-000794 compared to MOD(OMe)-scrambled.

**Table 2.**
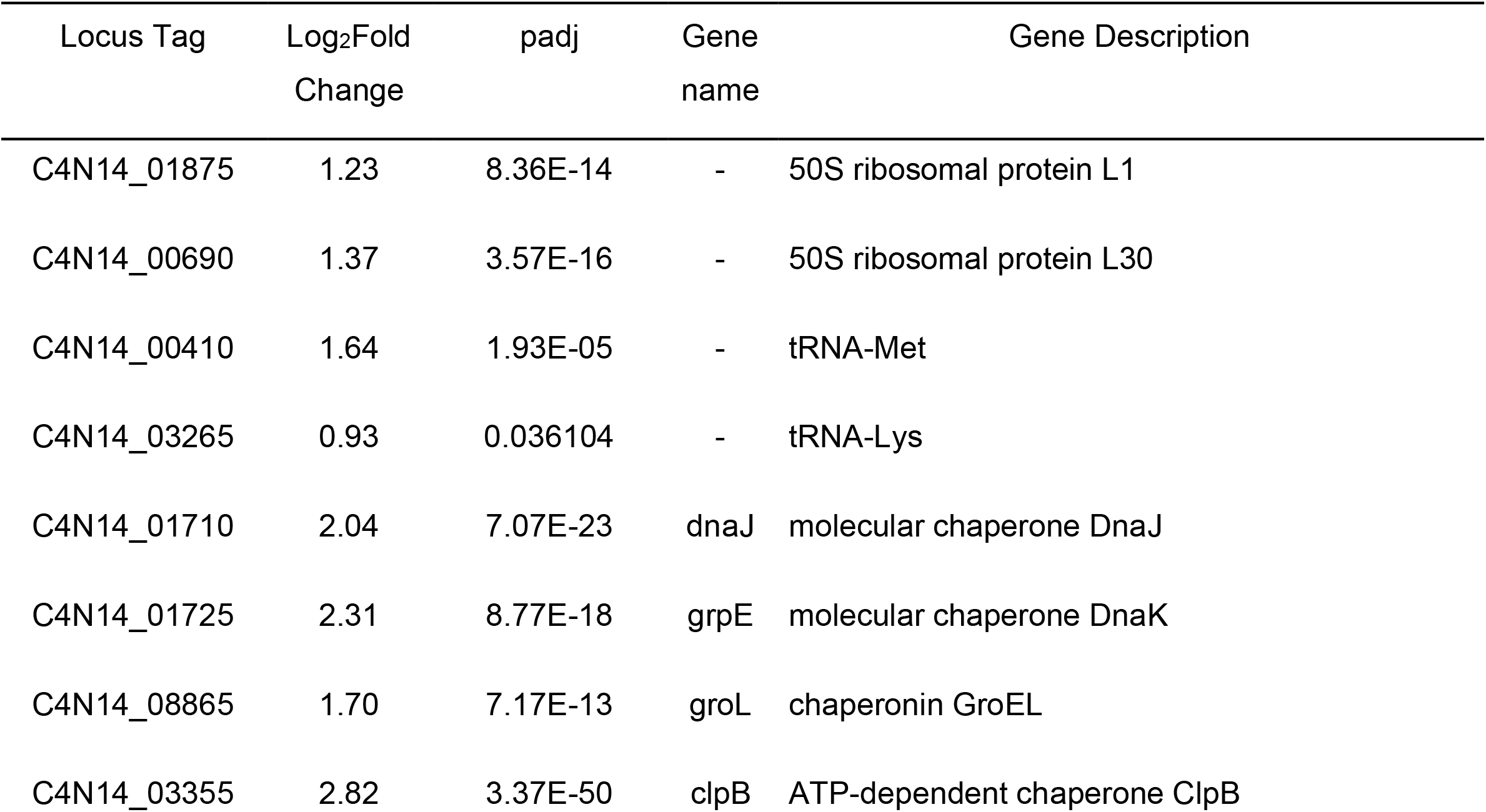
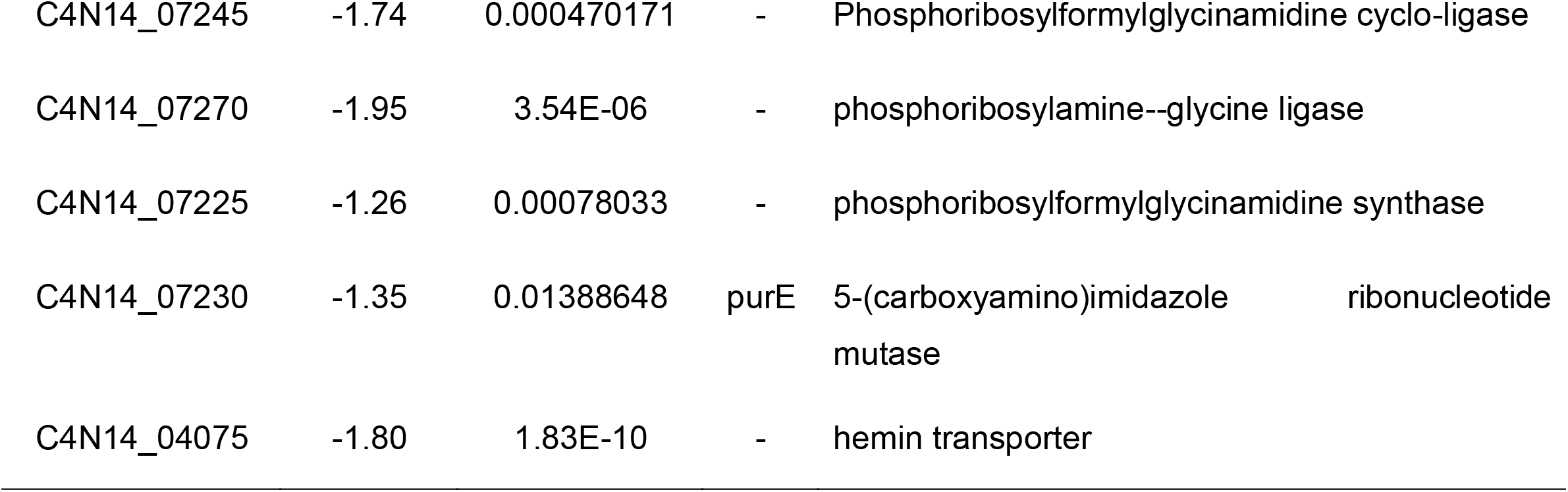
Differentially expressed genes after 5 h treatment with 500 nM MOD(OMe)- 000794 (Relative to MOD(OMe)-scrambled RNA control).

**Fig. 4.**
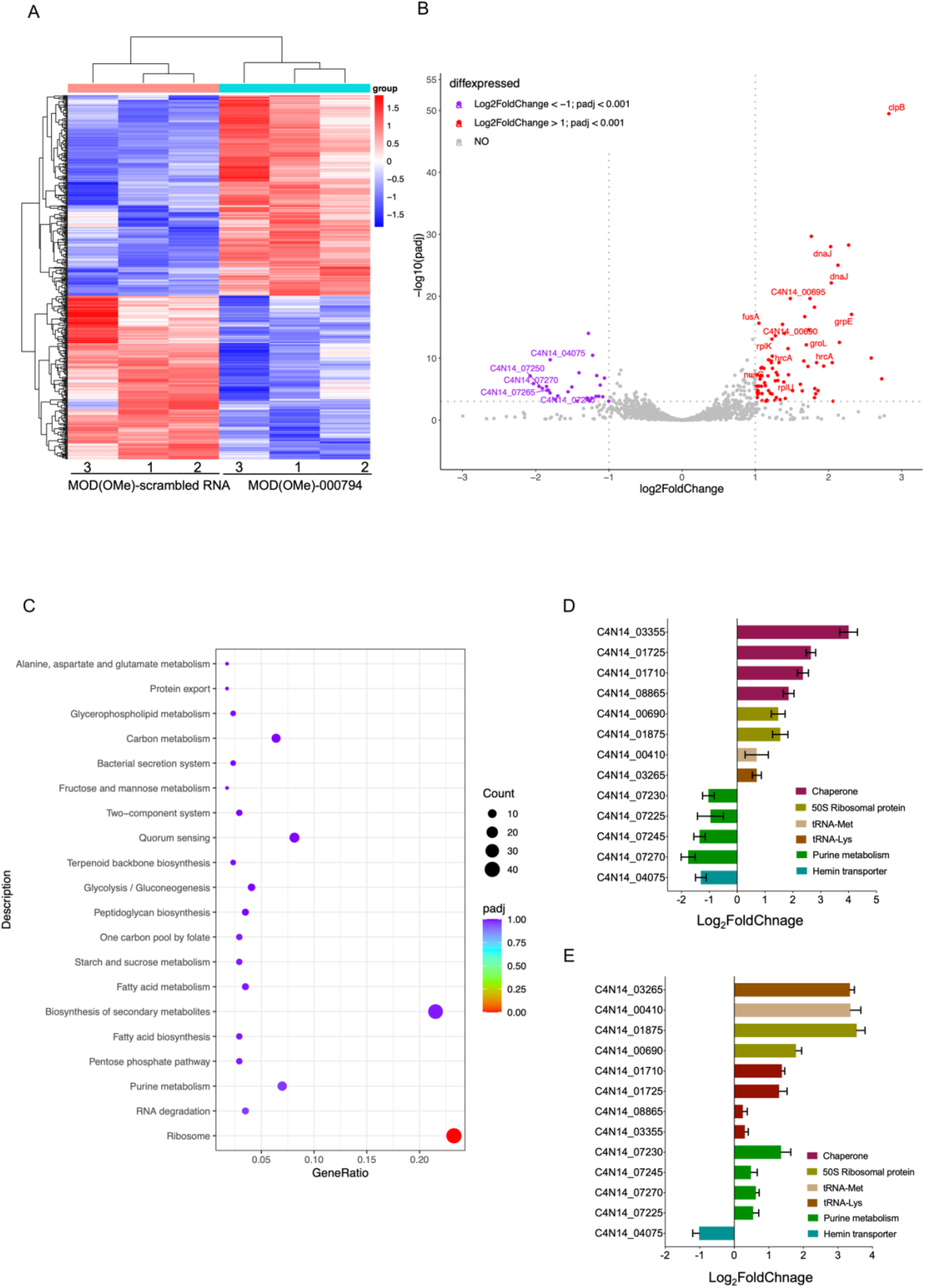
(*A*) Cluster analysis showing differentially expressed genes upon the indicated treatment conditions. Each comparison includes triplicate RNA-seq samples for the indicated MOD(OMe)-000794 or MOD(OMe)-scrambled. The coloring indicates log_2_FoldChange (FC) of the selected samples, while red and blue denote up- and down- regulation, respectively. (*B*) Volcano plots showing transcriptomic changes of *Fn* ATCC 23726 in response to 500 nM MOD(OMe)-000794 relative to the MOD(OMe)-scrambled control RNA at 5 h. Shown in the plot are false discovery rate (FDR)-adjusted P-value (– log_10_, y-axis) and fold change (log2, x-axis). Significantly differentially regulated genes are characterized by an absolute fold change >2 (down-regulated log2 < −1, up-regulated log_2_ > 1; vertical dashed line) and an FDR-adjusted P-value < 0.001 (–log_10_ >3, horizontal dashed line). A full list of differentially expressed genes can be found in supplemental excels. (*C*) KEGG enrichment scatter plot of differentially expressed genes. The y-axis shows the name of the pathway, and the x-axis shows the Rich factor. Dot size represents the number of different genes, and the color indicates the q-value. (*D*) Validation of differentially expressed genes during MOD(OMe)-000794 treatment. RNAseq results were validated by qRT-PCR. Relative gene expression (log_2_FoldChange) was normalized to 16S reference gene by the 2^−ΔΔCt^ method, relative to the MOD(OMe)-scrambled control. Values are mean ± SEM of three biological replicates. (*E*) Validation of differentially expressed genes during 1 μg mL^-1^ thiamphenicol treatment.

The upregulation of chaperons and ribosomal proteins by MOD(OMe)-000794 relative to MOD(OMe)-scrambled appears to be counterintuitive since we previously showed that tsRNA-000794 resulted in a global translation attenuation in *Fn* ATCC 23726 using a click chemistry labeled amino acid^[14]^. However, it has been reported that expression of certain ribosomal proteins increased when bacteria were challenged with ribosome-targeting antibiotics^[36, 37]^. For example, antibiotics targeting the 50S subunit of the ribosome (chloramphenicol or its analog thiamphenicol) or disruption of ribosome assembly via overexpression of a translational repressor ribosomal protein can upregulate the levels of ribosomal protein mRNAs, tRNAs and rRNAs in *E. coli*, suggesting a negative feedback mechanism to cope with translation suppression^[36, 37]^. Indeed, when treating *Fn* ATCC 23726 with thiamphenicol at a sub-minimal inhibitory concentration (1 μg ml^-1^), genes with >2-fold down- or up-regulation from the MOD(OMe)-000794 treatment group exhibited similar trends in the putative pathways associated with hemin, chaperon, ribosomal protein mRNAs and tRNA clusters, with the exception that the putative purine biosynthesis genes showed the opposite trend in gene expression under the MOD(OMe)- 000794- and thiamphenicol-treated conditions (Fig. 4*E*). Thus, we hypothesized that one of the mechanisms for tsRNA-mediated growth inhibition could be targeting ribosomes or protein synthesis such that impaired protein translation leads to mis-folding of proteins and subsequent recruitment of protein chaperons. Since the sequence of MOD(OMe)- 000794 matches part of the full tRNAs in *Fn*, it is plausible that MOD(OMe)-000794 acts as decoy tRNA to target the ribosome or rRNA and interfere with protein translation as reported previously^[38]^. To test the hypothesis, we biotinylated tsRNAs at either 5’ or 3’ and performed RNA affinity pulldown from the total cell lysate of *Fn* 23726. Several 30S and 50S ribosomal proteins were found to be enriched in biotinylated MOD(OMe)-000794 relative to that of biotinylated MOD(OMe)-scrambled (Dataset S1). While the pulldown assay cannot exclusively identify which exact ribosome subunit is directly targeted by MOD(OMe)-000794, our data indicate that MOD(OMe)-000794 induced a potent translational attenuation in *Fn*, likely through targeting ribosome components and triggered upregulation of corresponding genes to mitigate the stress.

To further probe how MOD(OMe)-000794 impacted *Fn* ATCC 23726 at the molecular and cellular levels, we adopted Raman spectroscopy that has become increasingly popular to measure different states of bacteria in a label-free fashion. The Raman spectrum represents an ensemble of molecular vibrations, providing comprehensive but complex data reflecting the metabolism and chemical compositions of the cells exposed to various drugs of different concentrations or durations^[39]^. To characterize the phenotypic differences between MOD(OMe)-000794 and MOD(OMe)-scrambled RNA control treatments in *Fn* ATCC 23726, we collected bacterial samples using the same treatment conditions that were used for the aforementioned bacterial RNAseq. Formaldehyde-fixed bacteria were subject to Raman spectra acquisition to measure different metabolic states and cellular compositions. As shown in Fig. 5*A*, several typical Raman peaks were identified corresponding to 720/780 cm^-1^ (DNA/RNA), 1003 cm^-1^ (phenylalanine), 1240 cm^-1^/1450 cm^-1^/1660 cm^-1^ (Amide I/II/III peaks), 2850 cm^-1^ (lipids and lipopolysaccharides), 2880 cm^-1^ (aliphatic amino acids). Through three-dimensional principal component analysis (PCA) of the Raman spectra, a global difference was detected between MOD(OMe)-000794 and MOD(OMe)-scrambled RNA control treated *Fn* ATCC 23726 (Fig. S14*A*). To understand which principal component contributes to the highest difference, we project the three-dimensional PCA into three two-dimensional PCA plots, respectively. As shown in Fig. S14*B-D*, PC3 contributes to the most difference between MOD(OMe)-000794 and MOD(OMe)-scrambled RNA control in *Fn* ATCC 23726. To identify which components from the PC3 underlined the phenotypic differences in the MOD(OMe)-000794 treatment group, we extracted the spectral information from PC3. Of note, peaks associated with the highest reduction from the MOD-tsRNA-00094 treatment are associated with proteins (containing aliphatic amino acids, 2880 cm^-1^) lipids^[40]^ and fatty acids (2850 cm^-1^) (Fig 5*B*)^[40]^. The reduction of lipids and fatty acids data agreed with the KEGG analysis from RNAseq, where the fatty acid metabolism represented the top five most enriched terms with gene downregulation (Fig. 4*C*). Additionally, downregulation of proteins containing aliphatic amino acids is in line with the putative ribosome-inhibiting roles of MOD-tsRNAs as shown in the RNAseq. Taken together, the data from Raman spectroscopy provided complementary evidence indicating the MOD-tsRNA-mediated interference of lipid metabolism and protein translation in *Fn*.

**Fig. 5.**
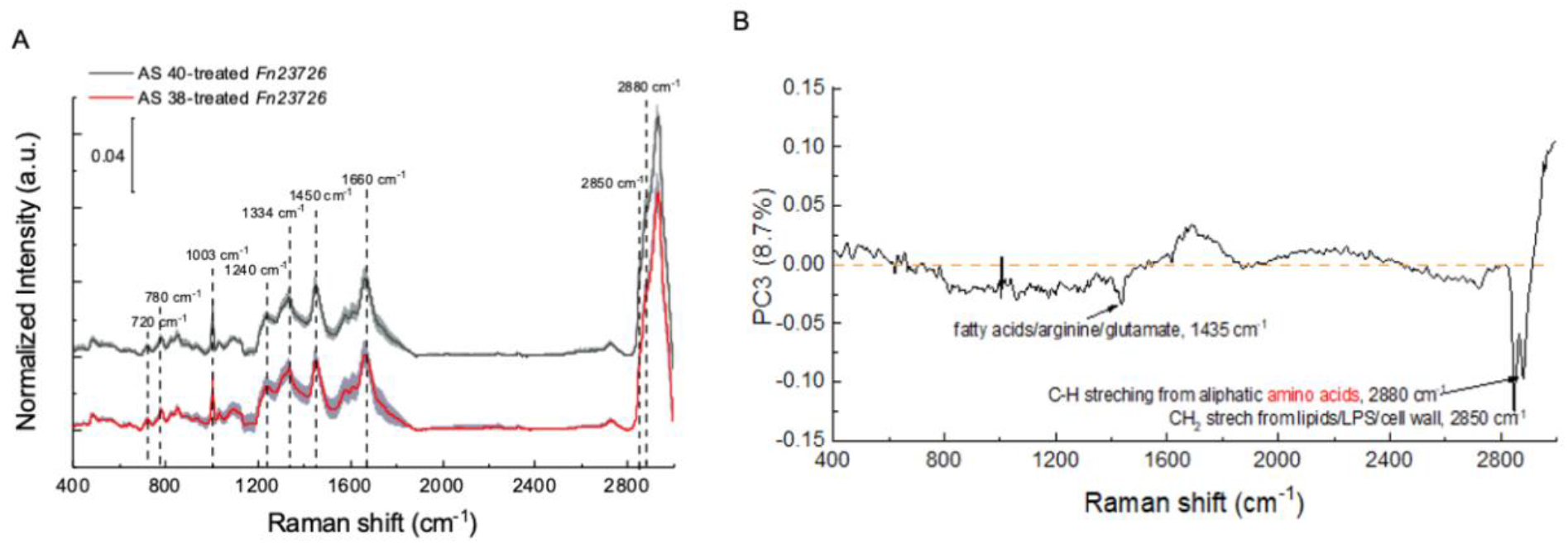
(*A*) Raman spectral signature reflects transcriptomic features of MOD-tsRNA treatment. Multiple Raman peaks showing up, 720/780 cm^-1^ (DNA/RNA), 1003 cm^-1^ (phenylalanine), 1240 cm^-1^/1450 cm^-1^/1660 cm^-1^ (Amide I/II/III peaks). (*B*) multiple chemicals such as proteins, especially proteins with aliphatic amino acids, glutamate, lipids illustrated the major composition of PC3, which suggests the metabolic difference of protein synthesis aroused between the treatments of MOD(OMe)-000794 (AS38) and MOD(OMe)-scrambled RNA (AS40).

## DISCUSSION

The chemical instability of sRNA molecules in general presents a formidable barrier to understanding the cross-kingdom functions of host sRNAs as well as developing RNA-based biologics for therapeutic applications. Over the last decades, chemically modified RNA nucleotides have greatly improved the nuclease resistance of RNAs while preserving their functionality. These efforts have propelled the development of nucleic acid-based genetic tools and therapeutics, including three recent FDA-approved small interfering RNA inhibitors for metabolic diseases, messenger RNA for vaccines, and CRISPR guide RNA for genome editing^[20-23]^. Building on the success of existing RNA technologies, our work here establishes tsRNAs as a new class of small RNAs for potential fundamental and applied studies in the context of host-microbe interactions. Specifically, we incorporated 2’-O-methylation and phosphorothioate chemistry at three terminal nucleotides of two host-derived tsRNAs, which were previously reported to play a role in cross-kingdom interactions between the host and *Fn*. Notably, such chemical modification strategy resulted in approximately 1000-fold higher potency than that of naturally occurring tsRNAs in inhibiting the growth of *Fn*. We found that the degree of modifications is critical to achieve the observed inhibitory efficacy as fully modified nucleotides abolished the inhibitory activities of tsRNAs. In addition to *Fn* type strains, the two MOD-tsRNAs also inhibited the growth of *Fn* clinical tumor isolates harvested from colorectal cancer specimens. In contrast, same MOD-tsRNAs at 10-time higher concentrations did not affect the growth of two representative oral bacteria, *Sm* ATCC 6249 and *Pg* ATCC 33277. Furthermore, the uptake studies suggested the internalization of tsRNAs by *Fn* but not *Sm* ATCC 6249 or *Pg* ATCC 33277. The uptake process is likely energy-dependent as the ATPase inhibitor sodium azide markedly blocked the internalization of tsRNAs by *Fn*. Taken together, these findings underscore species- and sequence-dependent modulation of bacterial growth by MOD-tsRNAs. Given the increasing implications of *Fn* in periodontal diseases, preterm birth, and cancer development and chemoresistance^[17-19]^, the MOD-tsRNAs developed in this study may establish host tsRNA as a first-of-its kind antimicrobial agent via chemical modifications to specifically target *Fn*-associated diseases.

To dissect the mechanisms of inhibition, we performed bacterial RNAseq to profile gene expression changes in *Fn* comparing MOD(OMe)-000794 to the scrambled control RNA. Interestingly, the majority of upregulated genes are associated with ribosomal proteins, tRNAs and protein chaperones, which were further validated by quantitative PCR. Of note, the KEGG analysis indicated that the protein translation represents the most enriched pathways targeted by MOD(OMe)-000794. The seemingly paradoxical upregulation of ribosome proteins during growth inhibition has been well documented in the literature and represents a hallmark of bacterial response when subjected to ribosome-targeting antibiotic treatments^[36, 37, 41, 42]^. In addition to their well-known inhibition of translation by interfering ribosomal functions, many of the ribosome-targeting antibiotics can directly bind to 30S or 50S ribosomal subunit precursors and inhibit the ribosomal assembly. These will often lead to increased expression of ribosomal protein-encoding genes^[37, 41]^. We speculated that MOD-tsRNAs may function as a new class of ribosome-targeting antimicrobials as the same set of genes were also upregulated when *Fn* was treated with a ribosome-targeting antibiotic, thiamphenicol. The upregulation of ribosomal protein mRNAs, tRNAs and protein chaperons may reflect bacterial stress responses during global translation inhibition mediated by MOD(OMe)-000794. To further corroborate the hypothesis, we carried out a biotinylated tsRNA pulldown assay and found that MOD(OMe)-000794 indeed recognizes several ribosomal proteins compared to MOD- scrambled RNA control. In addition to targeting the translation machinery, putative genes related to purine synthesis and hemin uptake are key downregulated genes and pathways upon the challenge of MOD(OMe)-000794. Since purine and hemin are essential for bacterial DNA synthesis and anaerobic growth, it is plausible to speculate that MOD- tsRNAs directly or indirectly interfere with multiple cellular functions to inhibit the growth of *Fn*.

The present work has also led to many intriguing questions. First, while findings from the global RNAseq implicate translation-related genes and pathways in the growth inhibition of *Fn*, the direct targets of MOD-tsRNAs remain elusive. It is possible that MOD-tsRNAs may directly interfere with ribosomal proteins to attenuate global mRNA translation in *Fn* mainly for two reasons: (1) No direct sequence complementarity was detected between the two *Fn*-targeting tsRNAs and *Fn* ATCC 23726 RNAs, thus arguing against an antisense mechanism. However, the central 21 nucleotides of tsRNA-000794 and 020498 match with the sense sequences of *Fn* tRNAs, suggesting that tsRNAs may be mis- incorporated into the ribosome machinery during active translation as the “ decoys” of tRNAs. (2) Despite current understanding of tsRNAs originated from studies in eukaryotic cells, most, if not all, tsRNAs have been shown to directly associate with RNA-binding proteins to affect mRNA stability and translation. Indeed, our biotinylated tsRNA pulldown assay supports the recognition of ribosomal proteins by tsRNAs. Second, it remains unclear how the tsRNAs define the species specificity. While our current findings suggest that the specificity is at least in part determined by an active uptake mechanism for tsRNAs in *Fn*, a putative transporter machinery for host sRNAs is yet to be identified in *Fn*. Of note, efforts have been made towards the identification of putative RNA importer proteins that facilitate internalization of extracellular sRNAs for intercellular or cross-kingdom gene modulation, such as Systemic RNA Interference Deficiency-1 (SID-1) in *C. elegan*, and the nematode homolog protein SID-1 transmembrane family member 1 (SIDT1) in mammalian cells. To the best of our knowledge, however, no such importer protein has been found in bacteria. Lastly, in addition to the selective uptake of certain tsRNAs by different bacteria, it is also possible that the intracellular targets of certain tsRNAs are only present in some bacteria such as *Fn*, which can dictate the bacterial range of different host-derived sRNAs in the context of cross-kingdom interactions. While we have focused on two tsRNAs in this present work, future work can investigate the full spectrum of host-derived tsRNAs or other sRNA molecules to uncover the fundamental mechanisms and full potential of host sRNAs underlying the host-microbiota interactions.

In summary, our work highlights an opportunity to use chemically modified RNA nucleotides to understand and harness host-derived tsRNAs to target opportunistic pathobionts, thus paving the way for fundamental research and therapeutic intervention.

## Materials and Methods

### Chemicals

All chemicals and cell culture broth were purchased from Fisher Scientific International Inc. (Cambridge, MA, USA) unless otherwise noted, and were of the highest purity or analytical grade commercially available. DNA and RNA oligos were ordered from MiliporeSigma (St. Louis, MO, USA) and Integrated DNA Technologies (Coralville, IA, USA). All molecular cloning reagents including restriction enzymes, competent cells, and the Gibson assembly kit were purchased from New England Biolabs (Ipswich, MA, USA).

### Bacterial Strains and Growth Conditions

*Fusobacterium nucleatum subsp. nucleatum* ATCC 23726, 25586, 10953, *Streptococcus mitis* ATCC 6249, and *Porphyromonas gingivalis* ATCC 33277 were purchased from the American Type Culture Collection (Manassas, VA, USA). *F. nucleatum* colon isolates were general gifts of Dr. Garrett Wendy at the Harvard T.H. Chan School of Public Health. *F. nucleatum* strains and *P*.*gingivalis* were cultured in liquid Columbia broth (CB) or on CB agar plates containing 5% defibrinated sheep blood (Hemostat laboratories, Dixon, CA, USA), and incubated at 37°C in an anaerobic chamber (Sheldon Manufacturing, Cornelius, OR, USA) containing 5% H2, 10% CO2, 85% N2. *S. mitis* were cultured in Brain-Heart Infusion (BHI) broth. *E. coli* K-12 were cultured in lysogeny broth (LB) media and incubated at 37 °C under aerobic conditions.

### Growth Inhibition Assays

Chemically synthesized tsRNAs and scrambled RNA control were reconstituted in sterile 1xPBS to obtain 500 μM stock concentrations stored at - 80 °C. 10 μL different chemical modified tsRNA were seeded into 96-well plates with 2- time serially diluted concentrations ranging from 5120 nM to 2 nM. The bacteria were diluted to OD_600_=0.01, and 90 μL diluted bacteria were added into individual wells of 96- well plates. The final concentrations were in the range of 1 to 512 nM. 10 μL of sterile 1xPBS was mixed with 90 μL diluted bacteria as an untreated control. Plates were incubated anaerobically at 37 °C, and OD_600_ was measured with a microplate reader at 0 h and 48 h to measure the outgrowth. For the CFU assay, 500nM MOD(OMe)-tsRNA treated bacteria were washed to remove free tsRNAs at indicated times, and 10-time serially diluted to enumerate CFU. Each growth inhibition assay was performed in three technical replicates and three biological repeats.

### tsRNA Stability Test and qPCR Quantification

A mixture of three MOD-tsRNAs and naturally occurring tsRNAs were added to pre-reduced Columbia Broth. 100 μl sample was frozen immediately by liquid nitrogen as the start point (0 h) and stored at -80 °C for the next step. Samples were incubated at 37 °C for 3 h, 6 h, and overnight before snap freezing. The collected samples were treated with 20 μg ml^-1^ of proteinase K at 50 °C for 30 min and then 1 mM EDTA and PMSF were added to the samples and incubated at 95 °C for 10 min to inactivate proteinase K. tsRNAs were reverse transcribed to cDNA with a HiFiScript cDNA Synthesis Kit (CoWin BioSciences, Cambridge, MA, USA) by stem-loop primers as previously described. cDNA was amplified and quantified by a QuantStudio 3 Real-Time PCR System (ThermoFisher, Waltham, MA, USA).

### MTT Cell Proliferation Assay

The normal oral keratinocyte (NOKSI) cell line was a gift from Dr. Silvio Gutkind and tested negative for mycoplasma contamination. Cells were cultured with a defined keratinocyte serum-free medium. The in vitro cell toxicity test was performed using the MTT assay (MilliporeSigma). MTT stock solution was prepared in sterilized 1xPBS and used with 1:10 dilution. NOSKI cells were seeded in 96 well plates at a concentration of 5000 cells/well and incubated with 100 µL of MTT reagent for 3 h inside the cell incubator. The converted dye is solubilized with dimethyl sulfoxide at the end of the incubation period. Finally, the cell viability in each well was measured at a wavelength of 570 nm with background subtraction at 650 nm.

### Fluorescence Microscopy of Cy3-tsRNA Labeled *Fn* Strains

Cy3 labeled tsRNA was reconstituted in 1xTBS containing 0.1 mM EDTA for imaging. Overnight-grown *Fn* was diluted to an OD_600_ of 0.1 and treated with 500 nM of 3’ Cy3 labeled tsRNA-000794, tsRNA-020498, and scrambled RNA control overnight (15-20 h) at an anaerobic chamber. Labeled bacteria were then washed with 0.9% NaCl for three times under a centrifuge speed of 17,000 x *g* for 10 min. Washed samples were then sandwiched between a cover glass and poly-L-lysine coated cover slide. Samples were then immediately imaged by a ZEISS LSM 800 confocal microscope with a fast Airyscan detector (with 120 nm lateral resolution and 350 nm axial resolution). To ensure the image quality, we utilized a 63x Plan-Apochromat NA1.4 oil immersion objective. Samples were excited at a wavelength of 514 nm with a 10% power and detected in the range of 550-600 nm. To quantify the fluorescence intensity from the same sample patch, dynamic range was adjusted to be the same under a channel-mode confocal modality. To have a clear visualization of tsRNA-Cy3 incorporation at subcellular level, super-resolution by Airyscan was achieved at a gain of 800 V. Images were visualized and analyzed by FiJi (NIH) and OriginLab (OriginLab Corporation).

### Fluorescence Microscopy of tsRNA-Alexa 488 Labeled *Fn* Strains

3’ Alex488 labeled tsRNA were reconstituted in 1xPBS. Overnight-grown *Fn* was diluted to an OD_600_ of 0.1 and treated with 250 nM tsRNA-000794-Alexa 488, tsRNA-020498-Alexa 488, and scrambled RNA-Alexa 488 control for 24h at an anaerobic chamber followed by three washes with 0.9% NaCl. Washed samples were then sandwiched between a cover glass and a SUPERFROST® PLUS-Adhesion Slide (Electron Microscopy Sciences, Hatfield, PA, USA). Samples were then immediately imaged by a Nikon, Epi-fluorescence microscope.

### Confocal Fluorescence Imaging of Cy3-tsRNA Uptake

Briefly, *Fn* ATCC 23726 with and without Cy3-tagged tsRNA treatment were centrifuged at a speed of 17,000 × *g* for 5 min, then washed with 1×PBS twice before putting the fresh samples on ice prior to the image acquisition. *Fn* was sandwiched between a cover slide and a #1.5 cover glass. Samples were excited at a wavelength of 514 nm and collected by a photomultiplier tube with an emission filter from 530 nm-600 nm. An oil-immersion 63× (NA=1.4) objective (Zeiss Objective Plan-Apochromat) was utilized to image samples.

### SYTOX Green Assay

*Fn* ATCC 23726 after treatment with MOD(OMe)-000794, MOD(OMe)-020498, and MOD(OMe)-scrambled RNA control was centrifuged at a speed of 17,000 × *g* for 5 min, then washed with 1×PBS. SYTOX green (S7020, ThermoFisher) was added to *Fn* at a working concentration of 5 μM in 1×PBS. Fluorescence intensity was then acquired after incubation at room temperature for 30 min. SYTOX green was excited at a wavelength of 488 nm and collected through a bandpass filter from 500-550 nm. An oil-immersion 63× (NA=1.4) objective (Zeiss Objective Plan-Apochromat) was utilized to image samples. Transmission images were acquired through a differential interference contrast setting. All the images were acquired under the same image acquisition setting and analyzed by FiJi and R. To quantify the fluorescence intensity from MOD(OMe)-000794 and MOD(OMe)-scrambled RNA control-treated *Fn*, integrated fluorescence intensity from the whole image was normalized by the total area of *Fn*. The higher the value indicates lower viability.

### Raman Spectroscopy

Cells were dissociated into single cells and fixed in 4% (V/V) paraformaldehyde for 15 min and were washed three times with 1xPBS. Before Raman measurements, the fixed cells were dropped onto an aluminum-coated Raman substrate to be air dried. Raman spectra were acquired using an HR Evolution confocal Raman microscope (Horiba Jobin-Yvon) equipped with a 532-nm neodymium-yttrium aluminum garnet laser. The laser power on cells was 12 mW after attenuation by neutral density filters. An objective with a magnification of 100× was used to focus single cells with a laser spot size of ∼1 μm^2^, and Raman scattering was detected by a charge-coupled device cooled at -70 °C. The spectra were acquired in the range of 400 cm^-1^ to 3,000 cm^-1^ with a 600 grooves per mm diffraction grating. A mapping mode was used to characterize single cells pooled from three biological replicates, and the acquisition parameters were 20 s per spectrum by averaging two times, around 20 spectra from three biological replicates. Each sample was performed in three biological replicates and three technical replicates. All SCRS were preprocessed by cosmic ray correction and polyline baseline fitting with LabSpec 6 (Horiba Scientific). Spectral normalization was done by vector normalization of the entire spectral region (normalized by the norm). The choice of vector normalization was made to correct general instrumentation fluctuation as well as sample and experimental variables (e.g., thickness of the sample) without strongly interfering with the nature of the biological content. Normalization using a particular component, such as nucleic acids or Amide I peak, was not used here, to avoid any presumptions of specific biomolecular changes. Final data were presented through R and OriginPro.

### Sodium Azide Treatment

Sodium azide was prepared at a stock concentration of 1 M in water. *Fn* ATCC 23726 with a starting OD_600_ of 0.1 were mixed with sodium azide at a concentration range of 0 to 10 mM. Moreover, tsRNA-Cy3 was added to the above scenarios at a working concentration of 500 nM. Samples were then incubated at anaerobic chamber overnight before taking out for image acquisition and OD_600_ measurement.

### RNA Isolation

Overnight bacteria culture was diluted to OD_600_ of 0.2 before the tsRNA treatment. 10 µL 100 µM MOD(OMe)-000794, MOD(OMe)-scrambled RNA, and 1xPBS were added to 2 ml diluted bacterial culture separately with three biological repeats at the 500 nM working concentration. Cell pellets were collected at 5 h at 13,000 x *g* 2min followed by snapping freeze in liquid nitrogen and stored at -80 °C until RNA extraction. RNA was extracted by RNApure Tissue & Cell Kit (CoWin Biosciences, CW0560S) as manufacturer’s instructions except for DNase I treatment. RNA was eluted with 30 µL 1x TE buffer (10 mM Tris-HCl, 0.1 mM EDTA, pH 7.5) twice followed by TURBO DNA-free kit (ThermoFisher, AM1907) treatment at 37 °C for 20 min. 5 µL of Dnase-treated RNA was used for reverse transcription later and the remainder RNA was stored at -80°C until RNA Sequencing. For thiamphenicol treatment group, overnight bacterial culture was diluted to OD_600_ of 0.5 followed by adding 6uL 1 mg mL^-1^ thiamphenicol or vehicle control (EtOH) to 6 mL diluted bacterial culture. After 5h treatment, collected the pellets and isolated RNA as description.

### RNA Sequencing and Data Analysis

RNA sample quality was assessed using a Nanodrop (ThermoFisher) and Agilent 5400 (Agilent Technologies, Santa Clara, CA, USA). Prokaryotic mRNA sequencing was performed using the NovaSeq PE150 platform at the Novogen facility (Sacramento, CA, USA). The library was prepared by a Ribo-Zero protocol (250-300 bp insert strand specific library with rRNA removal using NEB Ribo- Zero Magnetic Kit). Paired-end sequencing produced 150 bp reads, to a depth of ∼2G output per sample. Sequences were mapped to a reference genome, *Fusobacterium nucleatum* ATCC 23726 (GenBank accession: CP028109) using a Bowtie2 pipeline adjusted for paired-end sequencing. Differential gene expression was analyzed using the DEseq2 pipeline in Rstudio as previously described^[43]^. Sequence data was filtered by removing genes with zero reads. The false discovery rate (FDR) was set to 5% and genes with a log_2_FoldChange of >1 or < -1 and a p-adjusted value (padj) < 0.001 were considered significant. All reported data are representative of three biological replicates.

### Real-Time PCR

Primers were designed by NCBI Primer BLAST and synthesized by MiliporeSigma. cDNA was synthesized from total RNA by HiFiScript cDNA Synthesis Kit (CoWin Biosciences). qRT-PCR was performed in 96-well plates format using FastSYBR Low Rox (CoWin Biosciences) on Quantstudio 3 Real-Time PCR System (ThermoFisher) using the following protocol: 95 °C for 20 s, 40 cycles of 95 °C for 3 s, and 60°C for 30 s, and a final cycle of 95°C for 15 s, 60 °C for 1 min, and 95 °C for 15 s. Relative gene expression was calculated by 2^−ΔΔCt^ method using 16S RNA as the reference gene. qRT-PCR was performed with three technical repeats and three biological repeats. Primers used for qRT-PCR are shown in Table S1.

### Biotinylated tsRNA Affinity Pulldown from Bacterial Lysate and Mass Spectrometry

Dynabeads® M-270 Streptavidin beads (ThermoFisher, Catalog# 65305) and streptavidin agarose resin (G-Biosciences. St. Louis, MO. U.S.A) were washed according to the instructions and blocked by 1 mg mL^-1^ BSA at RT for 1 h. M-270 beads and resin were washed by 3xTBS (Tris HCl 150 mM, NaCl 0.45 M, ph7.4) and 1xTBS (Tris HCl 50 mM, NaCl 0.15 M, ph7.4) three times respectively and resuspended in the same volume buffer as the initial volume of beads taken from vial by 3xTBS and 1xTBS. 50 mL bacterial culture was harvest by spinning at 4000 rpm 15min followed by 3 times 1xPBS washing steps. Cell pellet was rotated at RT for 20 min in 3 mL cell lysis buffer (50 mM Tris HCl, 0.15 M NaCl, ph7.4, 1 mM DTT, 0.5%(V/V) NP-40, 50 μg mL^-1^ lysozyme and protease inhibitor cocktail EDTA free). Lysates were sonicated on ice with 5 s on and 5 s off for a total 5 min at 50% power (Soniprobe, Dawe Instruments, England) for 3-4 times until the lysate was transparent and cleared by centrifugation. Heparin was added to the clear extracts with 100 μg mL-1 working concentration followed by incubation with precoated resin at 4 °C for 30 min and quantified for protein concentration by DC Protein Assay reagent package. Before incubating with pretreated M-270 beads, 1 mM MgCl_2_, 1 mM ATP and 40U mL^-1^ RnaseOUT were added to the precleared cell lysates. 5’ or 3’ Biotin- tsRNA with the final concentrations 500 nM was added to 10 μL precoated M-270 beads and incubated on ice for 30mins. After 3 times of washing by 1xTBS, the precleared lysates were aliquoted to the pretreated M-270 beads equally and incubated at RT for 2 h on the rotator. M-270 beads were washed for 10 min at RT with wash buffer (50 mM Tris HCl, 0.15 M NaCl, pH 7.4, 0.5%(V/V) NP-40) twice followed by 1xTBST (25 mM Tris, 0.15 M NaCl, ph 7.4, 0.05%(V/V) Tween™-20) twice. Beads were sent to Poochon Scientific, CT, USA for mass spectrometry analysis.

### Statistical Analysis

All statistical analyses were performed using GraphPad Prism 9 (San Diego, CA, USA). Data were analyzed with one-way or two-way analysis of variance (ANOVA) followed by *Bonferroni* test for statistical significance.

## ACKNOWLEDGEMENTS

This work was supported by the Northeastern University Faculty start-up funding (JL), National Institute of Biomedical Imaging and Bioengineering (R21EB030769 (JL)), National Institute of Dental and Craniofacial Research (NIDCR) awards (R03DE031329 (JL), R01DE023810 (XH), R01R01DE030943 (XH)), and Forsyth Institute Collaborative Pilot grant FSI_CP05 (XH). *Fn* CTI isolates from CRC specimens were kindly provided by Dr. Wendy Garret from Harvard T.H. Chan School of Public Health. We are grateful to Dr. Susan E. Abbatiello, the director of the Barnett Institute of Chemical and Biological Analysis, Department of Chemistry and Chemical Biology at Northeastern University for scientific advice. We would like to express our gratitude to Dr. Chenggang Wu at UTHealth Houston’s McGovern Medical School, Dr. Daniel Slade at Virginia Tech and Dr. Yiping W. Han at Columbia University for the helpful discussions. We also wish to thank Li Lab members Xin Sun, Yun Ni and Shaobo Yang for their generous instructions for protein purification and stem loop qPCR primers design.

## References

1. Zheng, D., T. Liwinski, and E. Elinav, Interaction between microbiota and immunity in health and disease. Cell Res, 2020. 30(6): p. 492–506.

2. Gu, M., et al., Host innate and adaptive immunity shapes the gut microbiota biogeography. Microbiol Immunol, 2022. 66(6): p. 330–341.

3. Diamond, G., et al., The roles of antimicrobial peptides in innate host defense. Curr Pharm Des, 2009. 15(21): p. 2377–92.

4. Chehoud, C., et al., Complement modulates the cutaneous microbiome and inflammatory milieu. Proc Natl Acad Sci U S A, 2013. 110(37): p. 15061–6.

5. Sawa, T., et al., Immunoglobulin for Treating Bacterial Infections: One More Mechanism of Action. Antibodies (Basel), 2019. 8(4).

6. Zeng, J., et al., Cross-Kingdom Small RNAs Among Animals, Plants and Microbes. Cells, 2019. 8(4).

7. Cai, Q., et al., Cross-kingdom RNA trafficking and environmental RNAi-nature’s blueprint for modern crop protection strategies. Curr Opin Microbiol, 2018. 46: p. 58–64.

8. Teng, Y., et al., Plant-Derived Exosomal MicroRNAs Shape the Gut Microbiota. Cell Host Microbe, 2018. 24(5): p. 637-652.e8.

9. Lee, H.J., Microbe-Host Communication by Small RNAs in Extracellular Vesicles: Vehicles for Transkingdom RNA Transportation. Int J Mol Sci, 2019. 20(6).

10. Koeppen, K., et al., Let-7b-5p in vesicles secreted by human airway cells reduces biofilm formation and increases antibiotic sensitivity of P. aeruginosa. Proc Natl Acad Sci U S A, 2021. 118(28).

11. Liu, S., et al., The Host Shapes the Gut Microbiota via Fecal MicroRNA. Cell Host Microbe, 2016. 19(1): p. 32–43.

12. Liu, S., et al., Oral Administration of miR-30d from Feces of MS Patients Suppresses MS-like Symptoms in Mice by Expanding Akkermansia muciniphila. Cell Host Microbe, 2019. 26(6): p. 779-794.e8.

13. Cai, Q., et al., Plants send small RNAs in extracellular vesicles to fungal pathogen to silence virulence genes. Science, 2018. 360(6393): p. 1126–1129.

14. He, X., et al., Human tRNA-Derived Small RNAs Modulate Host-Oral Microbial Interactions. J Dent Res, 2018. 97(11): p. 1236–1243.

15. Li, S., Z. Xu, and J. Sheng, tRNA-Derived Small RNA: A Novel Regulatory Small Non-Coding RNA. Genes (Basel), 2018. 9(5).

16. Max, K.E.A., et al., Human plasma and serum extracellular small RNA reference profiles and their clinical utility. Proc Natl Acad Sci U S A, 2018. 115(23): p. E5334–e5343.

17. Brennan, C.A. and W.S. Garrett, Fusobacterium nucleatum - symbiont, opportunist and oncobacterium. Nat Rev Microbiol, 2019. 17(3): p. 156–166.

18. Han, Y.W., Fusobacterium nucleatum: a commensal-turned pathogen. Curr Opin Microbiol, 2015. 23: p. 141–7.

19. Yu, T., et al., Fusobacterium nucleatum Promotes Chemoresistance to Colorectal Cancer by Modulating Autophagy. Cell, 2017. 170(3): p. 548-563.e16.

20. Zhang, M.M., et al., The growth of siRNA-based therapeutics: Updated clinical studies. Biochem Pharmacol, 2021. 189: p. 114432.

21. Polack, F.P., et al., Safety and Efficacy of the BNT162b2 mRNA Covid-19 Vaccine. N Engl J Med, 2020. 383(27): p. 2603–2615.

22. Yin, H., et al., Structure-guided chemical modification of guide RNA enables potent non-viral in vivo genome editing. Nat Biotechnol, 2017. 35(12): p. 1179–1187.

23. Hendel, A., et al., Chemically modified guide RNAs enhance CRISPR-Cas genome editing in human primary cells. Nat Biotechnol, 2015. 33(9): p. 985–989.

24. Liang, H., et al., 3’-Terminal 2’-O-methylation of lung cancer miR-21-5p enhances its stability and association with Argonaute 2. Nucleic Acids Res, 2020. 48(13): p. 7027–7040.

25. Ohara, T., et al., The 3’ termini of mouse Piwi-interacting RNAs are 2’-O-methylated. Nat Struct Mol Biol, 2007. 14(4): p. 349–50.

26. Kirino, Y. and Z. Mourelatos, Mouse Piwi-interacting RNAs are 2’-O-methylated at their 3’ termini. Nat Struct Mol Biol, 2007. 14(4): p. 347–8.

27. Hassler, M.R., et al., Comparison of partially and fully chemically-modified siRNA in conjugate-mediated delivery in vivo. Nucleic Acids Res, 2018. 46(5): p. 2185–2196.

28. Kostic, A.D., et al., Genomic analysis identifies association of Fusobacterium with colorectal carcinoma. Genome Res, 2012. 22(2): p. 292–8.

29. Bullman, S., et al., Analysis of Fusobacterium persistence and antibiotic response in colorectal cancer. Science, 2017. 358(6369): p. 1443–1448.

30. Abed, J., et al., Fap2 Mediates Fusobacterium nucleatum Colorectal Adenocarcinoma Enrichment by Binding to Tumor-Expressed Gal-GalNAc. Cell Host Microbe, 2016. 20(2): p. 215–25.

31. Wu, X. and J.A. Hammer, ZEISS Airyscan: Optimizing Usage for Fast, Gentle, Super-Resolution Imaging. Methods Mol Biol, 2021. 2304: p. 111–130.

32. Han, Y.W., et al., Interactions between periodontal bacteria and human oral epithelial cells: Fusobacterium nucleatum adheres to and invades epithelial cells. Infect Immun, 2000. 68(6): p. 3140–6.

33. Popella, L., et al., Global RNA profiles show target selectivity and physiological effects of peptide-delivered antisense antibiotics. Nucleic Acids Res, 2021. 49(8): p. 4705–4724.

34. Rogers, A.H., et al., Aspects of the growth and metabolism of Fusobacterium nucleatum ATCC 10953 in continuous culture. Oral Microbiol Immunol, 1991. 6(4): p. 250–5.

35. Han, Y.W., Laboratory maintenance of fusobacteria. Curr Protoc Microbiol, 2006. Chapter 13: p. Unit 13A.1.

36. Dennis, P.P., Effects of chloramphenicol on the transcriptional activities of ribosomal RNA and ribosomal protein genes in Escherichia coli. J Mol Biol, 1976. 108(3): p. 535–46.

37. Takebe, Y., et al., Increased expression of ribosomal genes during inhibition of ribosome assembly in Escherichia coli. J Mol Biol, 1985. 184(1): p. 23–30.

38. Gebetsberger, J., et al., tRNA-derived fragments target the ribosome and function as regulatory non-coding RNA in Haloferax volcanii. Archaea (Vancouver, B.C.), 2012. 2012: p. 260909.

39. Germond, A., et al., Raman spectral signature reflects transcriptomic features of antibiotic resistance in Escherichia coli. Commun Biol, 2018. 1: p. 85.

40. Czamara, K., et al., Raman spectroscopy of lipids: a review. Journal of Raman Spectroscopy, 2015. 46(1): p. 4–20.

41. Champney, W.S., The other target for ribosomal antibiotics: inhibition of bacterial ribosomal subunit formation. Infect Disord Drug Targets, 2006. 6(4): p. 377–90.

42. Maitra, A. and K.A. Dill, Modeling the Overproduction of Ribosomes when Antibacterial Drugs Act on Cells. Biophys J, 2016. 110(3): p. 743–748.

43. Love, M., et al., RNA-Seq workflow: gene-level exploratory analysis and differential expression [version 2; peer review: 2 approved]. F1000Research, 2016. 4(1070).

