## Supplementary Figures for "Targeting *Fusobacterium nucleatum* through Chemical Modifications of Host-Derived Transfer RNA Fragments"

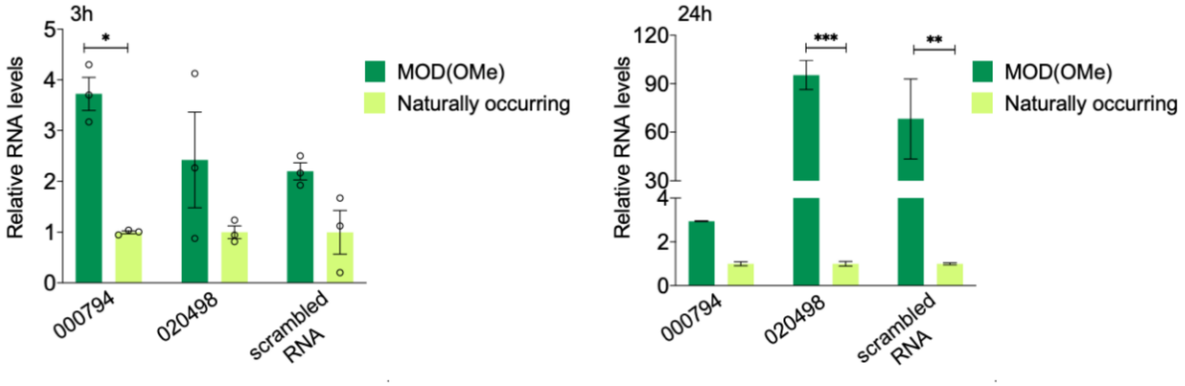


**Figure S1**: The enhanced stability of MOD-tsRNA over naturally occurring tsRNA at 3 and 24 h timepoints. The levels of intact MOD(OMe)-000794, MOD(OMe)-020498, MOD(OMe)-scrambled RNA, and corresponding mimics of naturally occurring RNAs were measured by the stem-loop reverse transcription PCR assay after incubation in the Columbia broth.

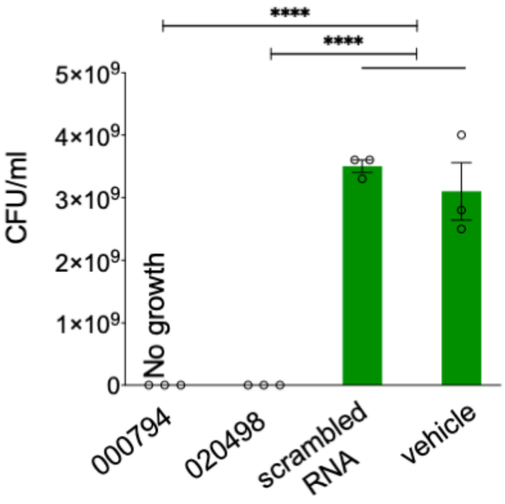


**Figure S2:** Quantification of CFUs after 24 h treatment with 500 nM MOD(OMe)-000794, MOD(OMe)-020498, MOD(OMe)-scrambled and vehicle control (1xPBS).

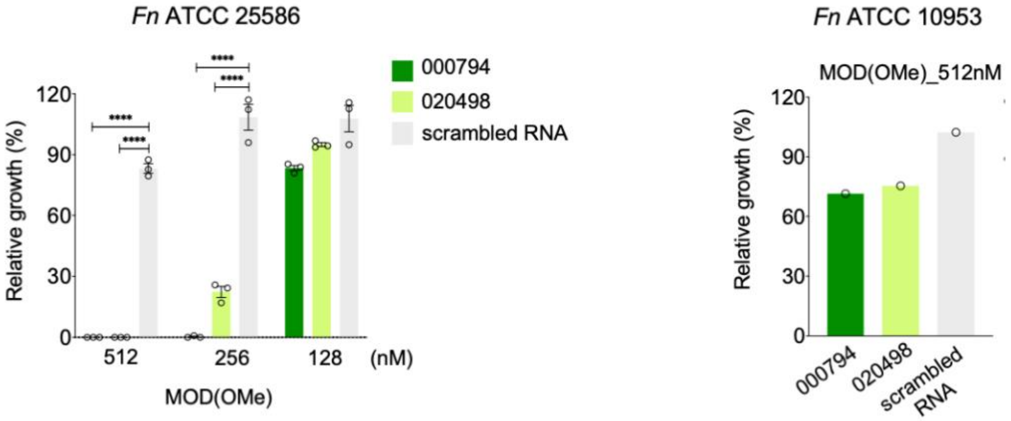


**Figure S3**: MOD(OMe)-000794 and MOD(OMe)-020498, but not MOD(OMe)-scrambled RNA, inhibited the growth of *Fn* ATCC 25586 and ATCC 10953. Higher concentrations of MOD(OMe)-tsRNAs were needed than those in *Fn* ATCC 23726 as shown in Fig 1*C*.

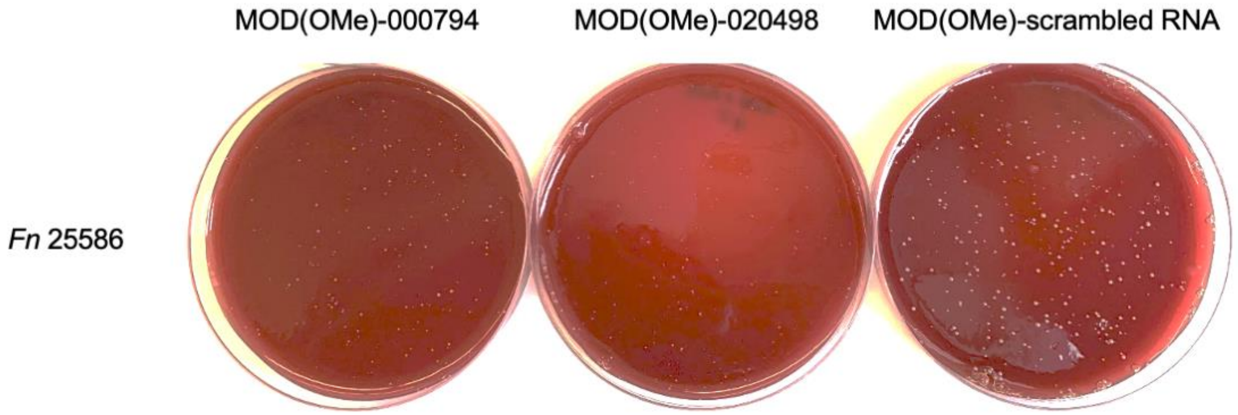


**Figure S4:** Representative images showing the CFU quantification for *Fn* ATCC 25586 following overnight treatment with 500 nM MOD(OMe)-tsRNAs in liquid culture under anaerobic conditions. Results are representative of two biological replicates.

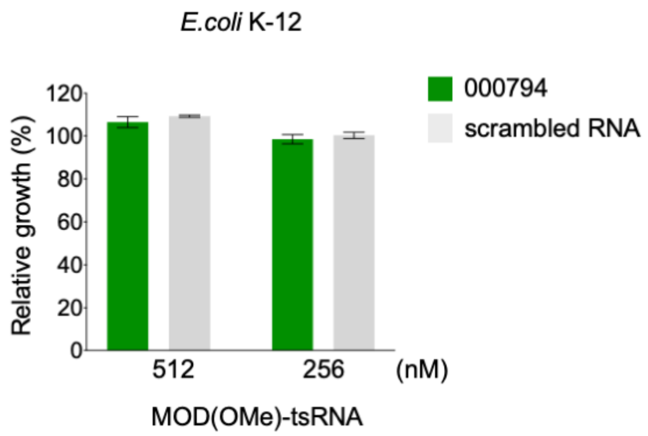


**Figure S5:** Lack of growth inhibition in *E.coli* K-12 by MOD(OMe)-000794 relative to MOD(OMe)-scrambled at 512nM.

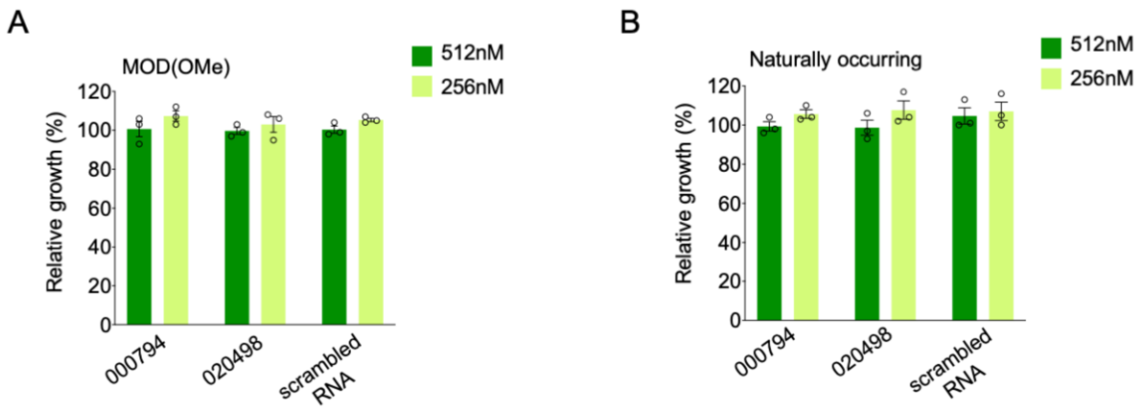


**Figure S6:** Relative growth rates of Normal Oral Keratinocytes-Spontaneously Immortalized (NOKSI) cells treated with MOD(OMe)-tsRNAs (*A*) or naturally occurring one (*B*) for 48 h. At 48 h, cell proliferation was measured by the MTT assay, and the growth rates were normalized to the untreated control groups (N=3). Growth data = Mean ± SEM, and are representative of two biological replicates.

A

| | |
|---|---|
| 000794 | [mC]*[mC]*[mG][mG][mC][mU][mA][mG][mC][mU][mC][mA][mG][mU][mC][mG][mG][mU][mA][mG][mA][mG][mC][mA][mU][mG][mA]*[mG]*[mA] |
| 020498 | [mG]*[mG]*[mG][mG][mG][mU][mA][mU][mA][mG][mC][mU][mC][mA][mG][mU][mG][mG][mG][mU][mA][mG][mA][mG][mC]*[mA]*[mU] |
| scrambled RNA | [mG]*[mG]*[mA][mC][mG][mA][mC][mA][mA][mG][mU][mU][mC][mG][mU][mG][mA][mC][mG][mA][mG][mC][mG][mC][mA][mU][mC]*[mU]*[mG] |

B

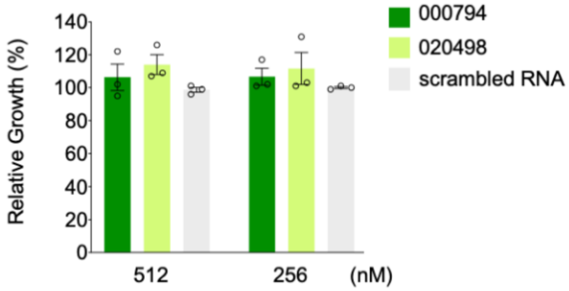


**Figure S7:** (*A*) Compositions and sequences of fully modified tsRNAs. * indicates a phosphorothioate (PS) bond, and [mA], [mU], [mC] and [mG] denote ribonucleotides with 2'-O-methylation. (*B*) Relative growth rates of *Fn* ATCC 23726 treated with fully modified tsRNAs. Full modifications of RNA backbone completely abolished the efficacy compared to partially modified tsRNAs (*i.e.*, MOD-tsRNAs shown in Figure 1). Growth data = Mean ± SEM, and are representative of three biological replicates.

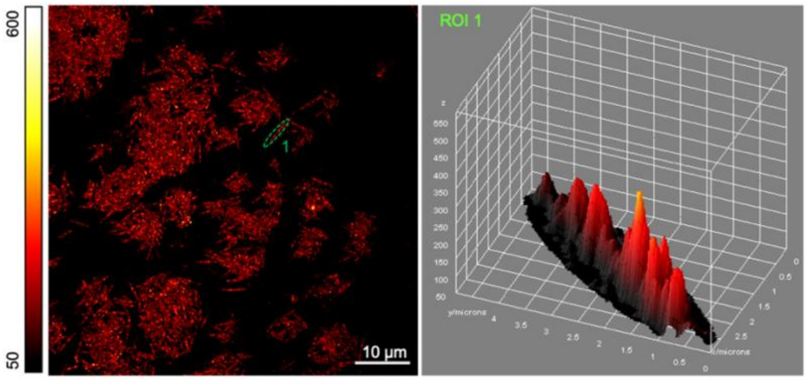


**Figure S8:** Internalization of tsRNA-000794-Cy3 by *Fn* ATCC 25586 imaged by Airyscan confocal microscopy.

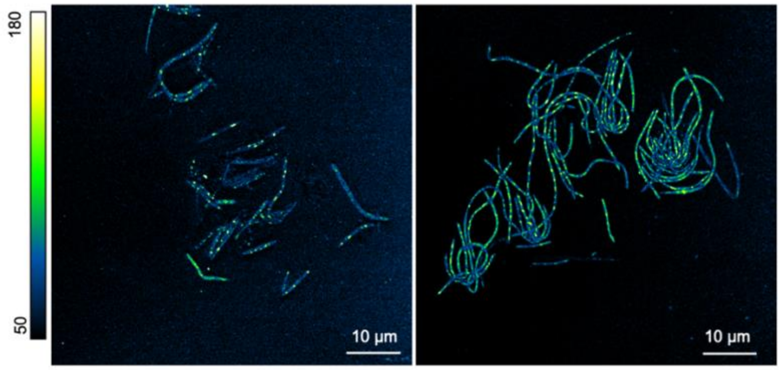


**Figure S9:** The uptake of tsRNA-000794-Cy3 by colon cancer-associated *Fn* isolates as imaged by Airyscan confocal microscopy.

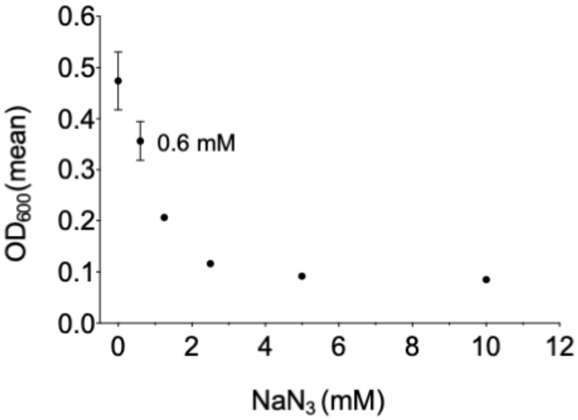


**Figure S10:** Titration of sodium azide concentrations in of *Fn* ATCC 23726. The growth of *Fn* was not significantly reduced at 0.6 mM. Bacterial culture at the log phase were diluted to OD600 of 0.1 before adding $NaN_3$ at a series indicated concentrations and incubated overnight under anaerobic conditions before $OD_{600}$ measurement.

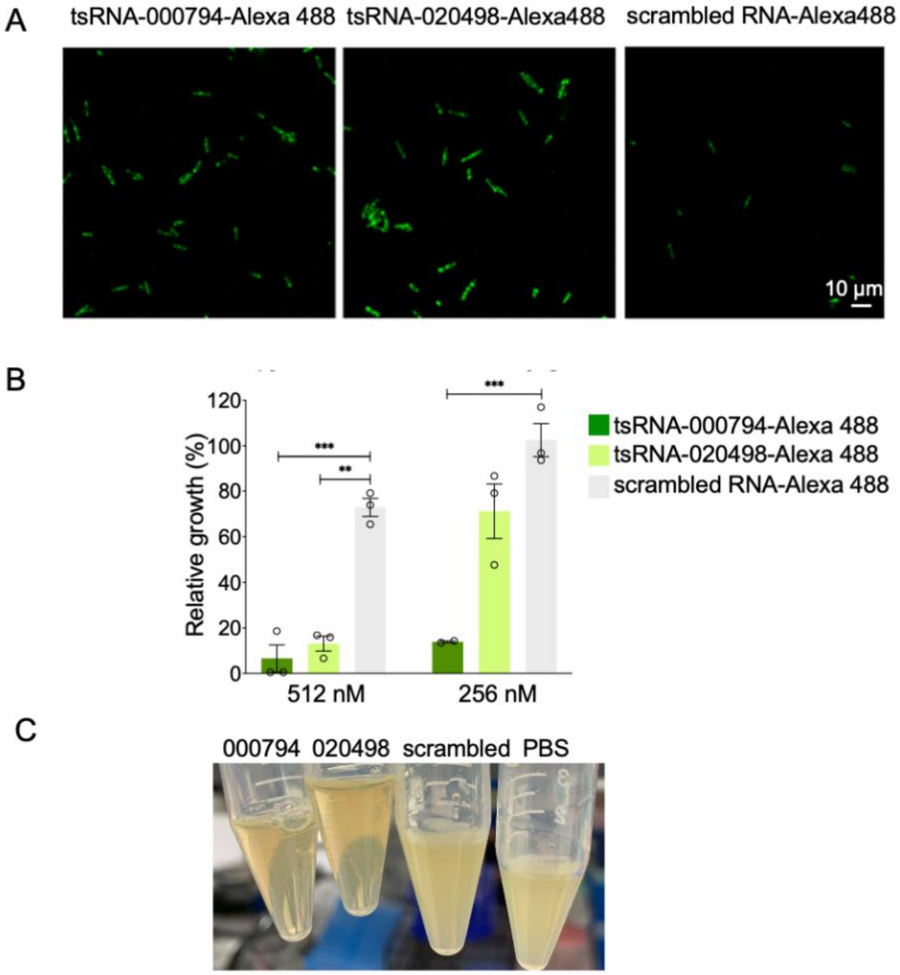


**Figure S11:** (*A*) Internalization of Alexa 488-labeled tsRNA-000794, tsRNA-020498 and scrambled RNA control in *Fn* ATCC 23726. Confocal microscopy demonstrated enhanced uptake of tsRNA-000794-Alexa 488 and tsRNA-020498-Alexa488 relative to the scrambled control-Alexa 488. (*B*) Growth inhibition of *Fn* ATCC 23726 by Alexa-488-conjugated tsRNA-000794 and tsRNA-020498 but not the scrambled control. Data are analyzed by the two-way ANOVA followed by Dunnett's Bonferroni multiple comparison tests. *$p < 0.05$, **$p < 0.01$, ***$p < 0.001$, ****$p < 0.0001$.(*C*) Representative images of *Fn* ATCC 23726 after overnight treatment with 512 nM of Alexa 488-labeled RNAs or PBS vehicle control.

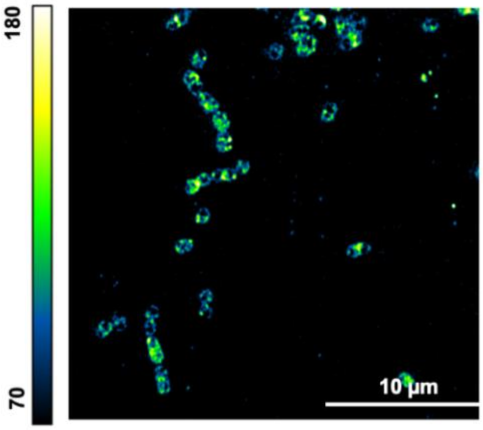


**Figure S12:** Periphery localization by tsRNA-000794-Cy3 of *S.mitis* ATCC 6249 through Airyscan confocal microscopy.

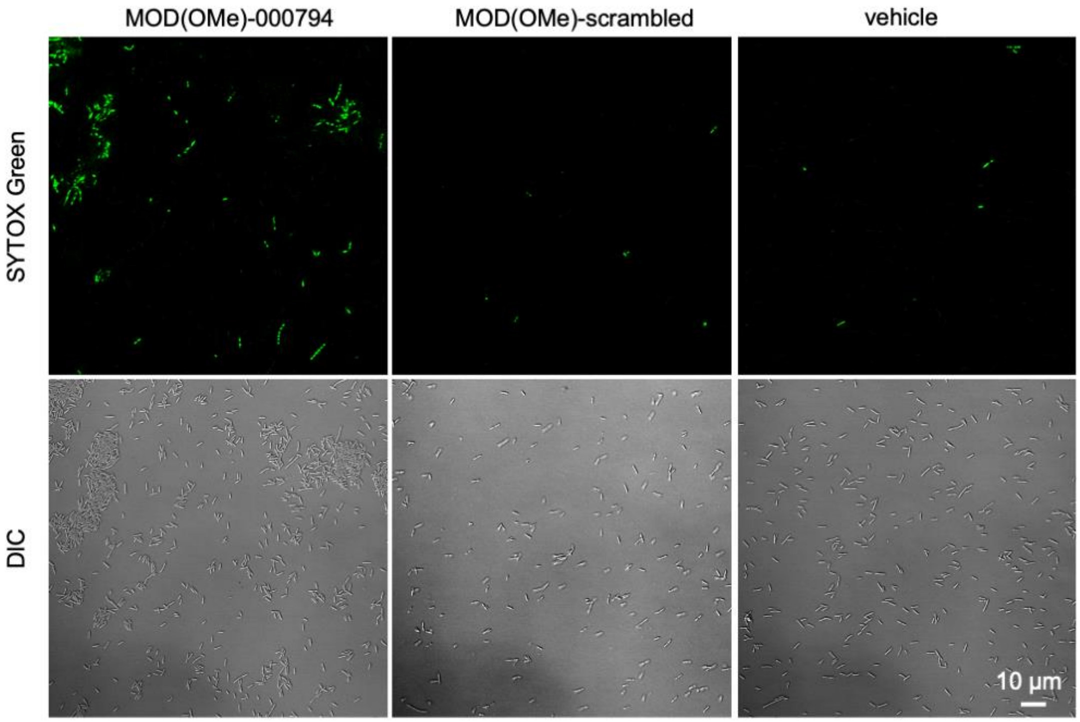


**Figure S13:** SYTOX green viability examination of *Fn* ATCC 23726 after treatment with 500 nM MOD(OMe)-000794, MOD(OMe)-scrambled RNA and 1xPBS buffer (vehicle control) for 5 h. Bacteria were treated at a starting $OD_{600}$ of 0.2. Images are representative of three biological replicates.

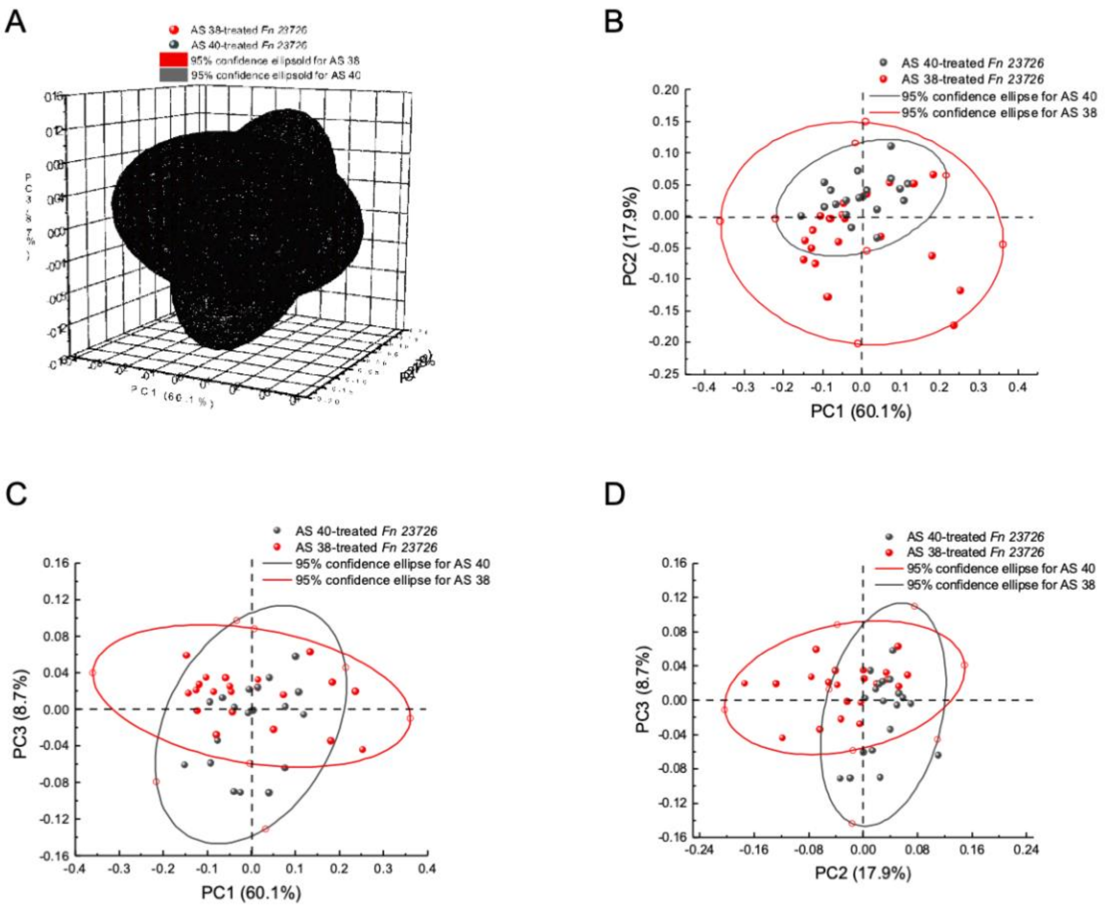


**Figure S14:** (*A*) Principal component analysis (PCA) of Raman spectra of *Fn* ATCC 23726 after treatment with 500 nM MOD(OMe)-000794 (AS38), MOD(OMe)-scrambled RNA (AS40) for 5 h. three-dimensional PCA plot indicates global difference as to the Raman spectra from MOD(OMe)-000794 and -scrambled RNA treated *Fn* ATCC 23726. Three-dimensional ellipse represents the 95% confidence interval. (*B*-*D*) three two-dimensional PCA plots suggests PC3 contributes to the most difference between MOD(OMe)-000794 (AS38) and -scrambled RNA (AS40) treated *Fn* ATCC 23726. Each dot represents a single Raman spectrum from *Fn* ATCC 23726 aggregates dried on the aluminum substrate. Two-dimensional ellipse represents the 95% confidence interval line.

**Table S1**. Primer sequences used for RNAseq verification.

| Primer name | Locus Tag | Sequence (5'-3') |
|---|---|---|
| Q176 | 16S | Fwd: CTTAGGAATGAGACAGAGATG |
| Q177 | | Rev: TGATGGTAACATACGAAAGG |
| Q273 | C4N14_01875 | Fwd: AGATCCAAGACATGCTGACCA |
| Q274 | | Rev: TCTGCTCCTGCAGCTAATGC |
| Q279 | C4N14_00410 | Fwd: GCTCAGCTGGATAGAGCAACGC |
| Q280 | | Rev: GGATCCAGCTGGACTCGAACCA |
| Q281 | C4N14_01710 | Fwd: TGGTGGCTTCAATGCAGGAG |
| Q282 | | Rev: AAACCTCCAAAGCCTCCACC |
| Q283 | C4N14_07245 | Fwd: AACTGCTGAAATGCCAGGCT |
| Q284 | | Rev: ATCCACTTGAAGCCACTGCT |
| Q289 | C4N14_07270 | Fwd: GAGAATGGGCGACCCTGAAA |
| Q290 | | Rev: ATCCACCTGCTGCCATAACC |
| Q291 | | Fwd: CGGCAGGAAAGGGTGTTGTA |
| Q292 | | Rev: TCTCCTGCAGCAGCAAATACT |
| Q297 | C4N14_07225 | Fwd: GCAGCACCTGTGGAAAATGT |
| Q298 | | Rev: GAACCAGTTGCTCCTCCACA |
| Q299 | C4N14_07230 | Fwd: AAGGGAGCAGCAGACTGTTT |
| Q300 | | Rev: AATGTGCTGCAAGTCCTGCT |
| Q301 | C4N14_01725 | Fwd: GCTGCTGCTCTTGCTTATGG |
| Q302 | | Rev: AATGTTCCCCCACCAAGGTC |

| | | |
|---|---|---|
| Q303 | | Fwd: CCAGCAGTTCAAGAATGGGT |
| Q304 | | Rev: CTTGTATTGCAGCACCTGCC |
| Q305 | C4N14_08865 | Fwd: TGAAAATATGGGGGCAGCCTT |
| Q306 | | Rev: TGTTGTTCCGTCTCCTGCAA |
| Q313 | C4N14_03355 | Fwd: GCCGTTAAATCCGTTGCTGA |
| Q314 | | Rev: TTACCAACCCCAGTAGGTCC |
| Q315 | C4N14_03265 | Fwd: CGACCATTAGAGTGCGGGAA |
| Q316 | | Rev: CGACAACCAGTGCCACAAAC |
| Q319 | C4N14_00690 | Fwd: TGCAGGTAAATGGGGAGCAA |
| Q320 | | Rev: GCAGAACCAGCTATTACCCCA |
| Q323 | C4N14_04075 | Fwd: TGATGCCTTAAGAGCGAAAGGT |
| Q324 | | Rev: TGGTCTCTATTTGCACTTGCT |
| Q325 | | Fwd: TGCATTAGGACATGAGTTTGGACA |
| Q326 | | Rev: TGCTGCTAAGGTATCTTCTGTTGC |

**Table S2**. Primer sequences used for MOD-tsRNA stability test.

| | | |
|---|---|---|
| Q169 | tsRNA-020498 stem loop | GTCGTATCCAGTGCAGGGTCCGAGGTATTCGCA CTGGATACGACATGCTCTA |
| Q189 | tsRNA-000794 stem loop | GTCGTATCCAGTGCAGGGTCCGAGGTATTCGCA CTGGATACGACTCTCATGC |
| Q190 | Scrambled RNA control stem loop | GTCGTATCCAGTGCAGGGTCCGAGGTATTCGCA CTGGATACGACcAGATGCG |
